# The shared genomic history of Middle to Late Holocene Southern Cone populations

**DOI:** 10.1101/2025.11.04.686483

**Authors:** Kim-Louise Krettek, Maria Barbara Postillone, Lucia Spangenberg, Javier Maravall López, Nicola Rambaldi Migliore, Ana Maria Chero Osorio, Hugo Naya, Ella Reiter, Tatiana Tondini, Mariano Bonomo, Valeria Bernal, Mariela E. Gonzalez, Nahuel Scheifler, Pablo G. Messineo, Gustavo Flensborg, Cristina Dejean, Alessandro Achilli, David Reich, Jose Lopez Mazz, S. Ivan Perez, Gustavo Politis, Gustavo Martínez, Cosimo Posth

## Abstract

The Southern Cone represents the southernmost region of South America to be colonized by humans. Although ancient genomes have been sequenced from southern Patagonia, genomic data from the central Southern Cone remain temporally and spatially sparse. The archaeological record of this region documents major cultural transformations during the Middle and Late Holocene, yet their relationship to demographic processes has long been debated. Here, we present genome-wide data from 52 individuals spanning the past 5,000 years, originating from four regions of the central Southern Cone in present-day Argentina and Uruguay: the central and southern Pampas, Northwest Patagonia, the Paraná River Delta and Lower Uruguay River, and the eastern lowlands of Uruguay. Genomic evidence from the Pampas reveals the presence of at least three distinct ancestries during the Middle Holocene. While genetic contacts with southern Patagonian groups were sporadic, we identify the expansion of an ancestry of unknown geographic origin by 4,800 years ago, which increased substantially during the Late Holocene. This same ancestry arrived in Northwest Patagonia by at least 600 years ago, and it co-existed with individuals carrying a southern Andean genetic profile until colonial times. Genetic structure differentiates populations along the Paraná River Delta and the Lower Uruguay River by approximately 1,600 years ago. In contrast, individuals from the eastern lowlands of Uruguay show genetic links with Sambaqui-associated populations from the southern coast of Brazil, suggesting the role of human dispersals in connecting tropical lowland cultural traditions. Overall, our work documents the diffusion of genetically distinct groups across all regions studied and provides compelling evidence that large-scale human movements contributed to the remarkable cultural diversity of central Southern Cone populations during the Middle and Late Holocene.

## Introduction

Within the last years, studies of ancient and modern genomes from Indigenous populations have shaped our understanding of the settlement history of South America at a sub-continental levels (Llamas *et al*., 2016; Moreno-Mayar *et al*., 2018; Posth *et al*., 2018; Arango-Isaza *et al*., 2023; Capodiferro *et al*., 2023; Gusareva *et al*., 2025). Once ancestral populations reached South America, a rapid genetic differentiation in multiple regions took place. At the same time, ancient genomic temporal series in different areas of South America have started to appear, including from the Bogotá Altiplano (Krettek *et al*., 2025), the Andes (Lindo *et al*., 2018; Nakatsuka *et al*., 2020a), southern Patagonia (Nakatsuka *et al*., 2020b; de la Fuente Castro *et al*., 2024), and the southern coast of Brazil (Ferraz *et al*., 2023). While in some areas large-scale replacements have been observed from the first forager groups until present-day Indigenous populations (Ferraz *et al*., 2023; Krettek *et al*., 2025), in others, long-lasting genetic continuity was traced back to the first settlers (Lindo *et al*., 2018; Nakatsuka *et al*., 2020a; Nakatsuka *et al*., 2020b). Understanding whether the distinct population dynamics observed in different regions are isolated cases or part of broader trends in certain areas of South America requires more geographically and temporally dense genomic sampling.

A region, which has been understudied in ancient human genomic work so far, is the Central Southern Cone (CSC) of South America. We define here CSC as the territories of Argentina, Uruguay, Chile, Paraguay, and southern Brazil between -25° and -40° latitude that comprises a large variety of ecologically and geologically different landscapes. Ancient genetic research in this vast region is mainly limited to mitochondrial DNA (mtDNA) studies (Crespo *et al*., 2018; Postillone *et al*., 2020a; Postillone *et al*., 2020b; García *et al*., 2021; Roca-Rada *et al*., 2021; Nores *et al*., 2022), with genome-wide data restricted to only 22 ancient individuals (Moreno-Mayar *et al*., 2018; Posth *et al*., 2018; Nakatsuka *et al*., 2020a; Nakatsuka *et al*., 2020b; Lindo *et al*., 2022; de la Fuente Castro *et al*., 2024). Since the beginning of the century, archaeological and bioanthropological models have been proposed to explain population dynamics and socio-environmental adaptations of hunter-gatherers from the final Late Pleistocene to the Late Holocene in CSC (Barrientos & Pérez, 2004; Bonomo, 2005; Politis *et al*., 2014; Perez *et al*., 2016; Martínez *et al*., 2017; Messineo *et al*., 2019; Politis & Borrero, 2024; among others). The dynamics of hunter-gatherer populations during the Middle Holocene (∼8,200-4,200 years before present (BP) particularly their spatio-temporal continuities, demographic decline, and population disruptions has been the subject of extensive debate. During this period, chronological and stratigraphic gaps in the archaeological record were recognized in both the Pampas (Martínez *et al*., 2023) and Northwest Patagonia (Barberena, 2015), in present-day Argentina. However, with increasing archaeological research and radiocarbon dating, these gaps have gradually been closing in both regions (Bonomo, 2013; Martínez *et al*., 2015; Timpson *et al*., 2021; Politis & Borrero, 2024). To explain these breaks in the record, archaeological and bioanthropological models referring to extinctions, expansions, bottlenecks, and population replacements among hunter-gatherer populations have been proposed for certain areas (Barrientos & Pérez, 2004; Barrientos & Perez, 2005; Barrientos, 2009; Barberena, 2015). However, other models support a different scenario of population and cultural continuity in the Pampas through time (Martínez *et al*., 2015; Mazzanti *et al*., 2015; Martínez *et al*., 2017; Martínez *et al*., 2024; Politis & Borrero, 2024). The retrieval of ancient genomic data from human remains dating to this period enables a direct assessment of the two competing hypotheses.

Moving forward in time, in the southern Pampas (often also described as the Pampa-Patagonia transition), a similar structure in the archaeological record and social and behavioural organizations of hunter-gatherers in the Mid-Holocene (up until 4,800 BP) was identified during the initial Late Holocene (from 3,100 BP). This chronological gap was interpreted as resulting from geological and taphonomic biases, and a scenario of cultural continuity was proposed (Martínez *et al*., 2024). In Northwest Patagonia, recent results have shown technological diversification and innovations with disparate spatio-temporal changes in the relative population density. An initial increase in population density during the Middle Holocene (by 6,500 BP) was followed by another increase in the Late Holocene (by 2,500 BP)(Cobos *et al*., 2022).

Finally, in the Late Holocene, hunter-gatherers from the Pampas underwent significant demographic and social transformations, also including technological adoptions and innovations (e.g., pottery, bow, and arrow), diversification and intensification in faunal resource exploitation, and complexity in funerary practices, thus suggesting the emergence of a new cultural organization (Politis, 2008). Specifically, around this period there is evidence of demographic growth, large camp-sites, increasing territoriality, technological changes, intensification processes, complexity in burials modalities, more competitive social interactions, construction of identities, and social differentiation, as well as a complex scenario of inter-ethnic contacts at local and extra regional levels (Barrientos & Pérez, 2004; Bonomo, 2005; Politis, 2008; Martínez *et al*., 2017; Beron, 2018; Messineo *et al*., 2023; Politis & Borrero, 2024). Similarly, in Northwest Patagonia, mobility patterns shifted during the Late Holocene when forager lifestyles were gradually replaced by more sedentary forms of residence as well as an increase in the differentiation of regional cultural developments (Perez *et al*., 2016; Cobos *et al*., 2022). Also, for the final phase of the Late Holocene, archaeological studies have debated scenarios of local population continuity versus replacements, but the available evidence has been insufficient for addressing this issue (Barrientos & Perez, 2005; Barberena *et al*., 2015; Martínez *et al*., 2015; Perez *et al*., 2016; Martínez *et al*., 2024).

Flowing into the La Plata River, the Paraná and Uruguay Rivers and their surrounding region have a longstanding archaeological tradition with evidence of fluvial adaptations, especially when considering the Middle Paraná-Salado and Paraná Delta (Gonzáles, 1977; Ceruti, 2003; Politis & Bonomo, 2023). The Late Holocene occupants of this region were hunter-gatherers/horticulturalists, known for their mound building and complex riverine settlements. The archaeological expression of these groups is known as the Goya-Malabrigo archaeological entity, characterized by elaborate ceramics and economy with hunting, fishing, gathering and small-scale horticulture (Bonomo *et al*., 2011; Politis & Bonomo, 2018). The Paraná Delta was also occupied by the Guaraní during the final phase of their expansion, approximately 700 years BP (Bonomo *et al*., 2015; Torino *et al*., 2023) but the settlement of the area by this group appears to have been relatively sparse.

In the region of present-day Uruguay, archaeological evidence throughout the Holocene suggests distinct population adaptation to riverine and wetland ecosystems with intensive use of terrestrial, lowlands, and coastal resources. The riverine adaptations of Uruguay River valleys share a very similar archaeological tradition with the Goya-Malabrigo of the Paraná River region (López Mazz *et al*., 2014; López Mazz, 2018; Politis & Bonomo, 2018). Mound-builders of the eastern lowlands of Uruguay were semi-sedentary hunter societies practicing horticulture (Bracco *et al*., 2000; López Mazz & Gianotti, 2001; Iriarte, 2006; Gianotti, 2015), and with mortuary traditions that suggest social differences between individuals (Gianotti & López Mazz, 2009). Importantly, the Late Holocene mounds and ceramics without decoration of eastern Uruguay provide hints for connections with tropical lowland cultures from southern Brazil.

A dynamic interaction between social, cultural, and biological factors is thus hypothesized across the CSC during the Middle and Late Holocene. To shed light on this genomically unexplored area, we generate new genome-wide and mtDNA data for 52 ancient individuals from four regions in Argentina and Uruguay, i.e., the central and southern Pampas, Northwest Patagonia, Paraná River Delta and Lower Uruguay River, and eastern lowlands of Uruguay, spanning a temporal transect from 5,200 to 200 BP. Our study explores the demographic processes of the CSC through time and investigates genetic diversity and similarities both within and between the regions to answer the following specific questions: 1) Determine the level of genetic continuity or discontinuity in populations from the Pampas and Northwest Patagonia during the Middle and Late Holocene; 2) Investigate the genetic similarities of ancient populations along the Paraná River Delta and Lower Uruguay River; 3) Estimate the genetic connections between pre-European individuals from the eastern lowlands of Uruguay and neighbouring regions. Our genomic findings are interpreted against the associated archaeological evidence to provide a more integrative understanding of the population history in each investigated region.

## Results

### Dataset and ancient DNA authentication

We generated genomic data from 52 individuals from 31 sites in four CSC regions: southern and central Pampas (11 sites, 21 individuals), Northwest Patagonia (13 sites, 15 individuals), Paraná River Delta and Lower Uruguay River (6 sites, 15 individuals), and Eastern lowlands of Uruguay (one site, one individual), dated to the Middle and Late Holocene (Figure 1, Table S1 and Supplementary Information). These genomic data were obtained from ancient DNA extracted from the petrous portion of temporal bones and from the crown of teeth, after generating double- and single-stranded libraries (see Materials and Methods). We captured the libraries in solution for a targeted set of ∼1.24 million single-nucleotide polymorphisms (1240K SNPs) and the entire mtDNA. We sequenced the libraries on Illumina platforms, obtaining an average coverage of ∼400K SNPs (12,515 to 968,033 SNPs). We then evaluated ancient DNA authenticity through mtDNA and X-chromosomal contamination estimates, which were found to be below 4% and 5%, respectively, except for three individuals (TPGU004, LAPE001, TPAG001), which were filtered for reads showing evidence of post-mortem damage (Table S1).

**Figure 1:**
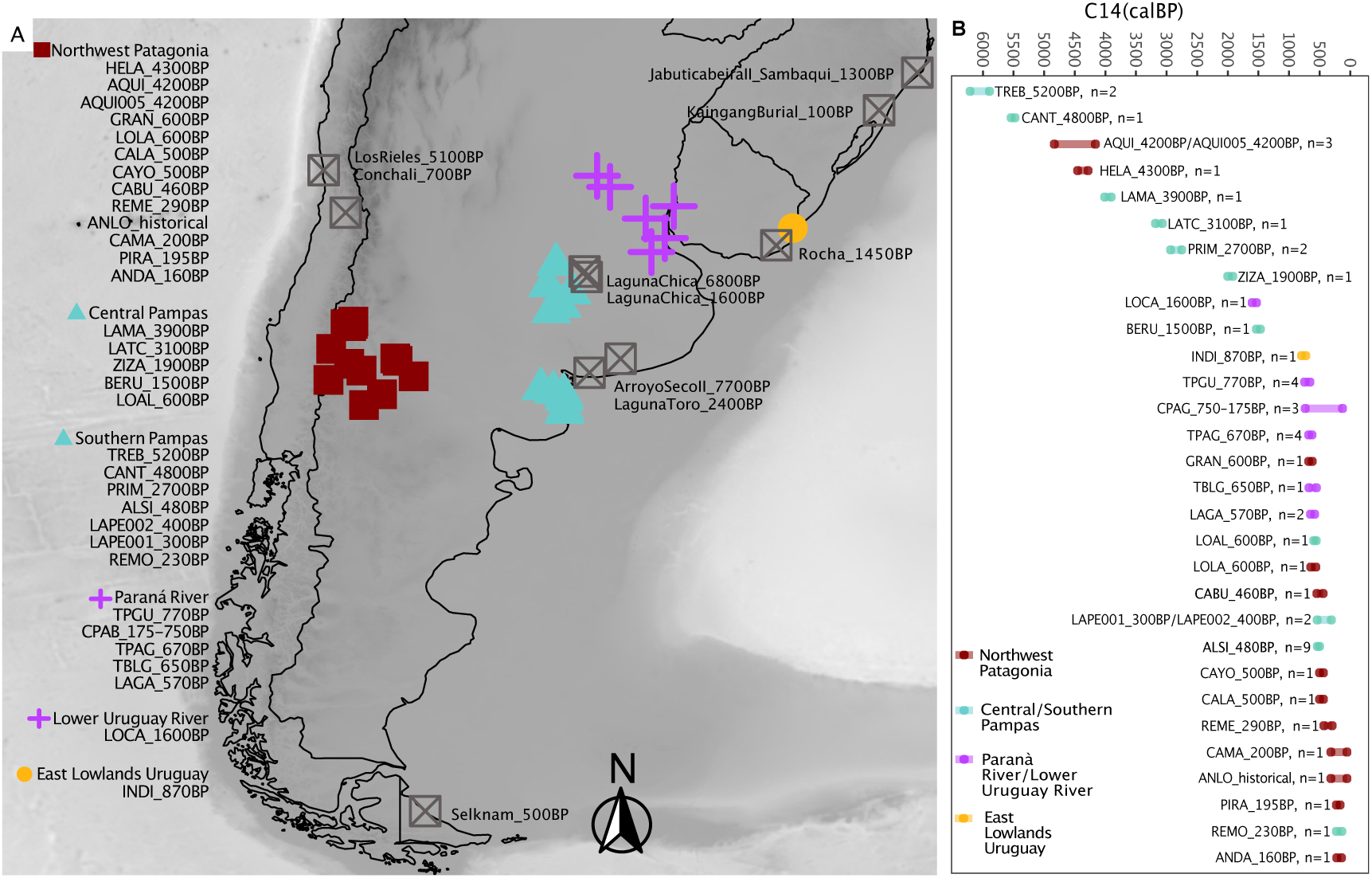
Geographical, temporal, and genetic overview of the newly generated dataset. A) Geographic map depicting all sites for which data is newly reported. Individuals have been grouped under one site label if genetic clustering has been confirmed (Material and Methods). B) Direct or associated radiocarbon dates for the analysed individuals from each archaeological site. When multiple dates per site were available, the full range after calibration is provided.

We performed a principal component analysis based on worldwide modern-day genomic allele-frequency variation (Mallick *et al*., 2016) and projected the genotypes of ancient individuals onto it. Confirming the absence of substantial allochthonous human DNA contamination, all ancient individuals fall within the variation of modern-day Indigenous Americans. The only exception is ALON001 from Northwest Patagonia, which is shifted toward present-day Europeans while lacking detectable levels of modern-day DNA contamination (Figure S1). These results, paired with its post-contact date, suggest that the genome of this individual carries recent European admixture.

### Uniparental markers and biological relatedness

Genetic sexing determined the presence of 27 females and 26 males among the studied individuals (Table S1). All male individuals belong to the Y-haplogroup Q1b1a and its subclades. This lineage is the predominant paternal haplogroup in Indigenous Americans, and our data confirm its high frequency in all analysed geographic regions (Huang *et al*., 2018; Grugni *et al*., 2019). The only exception is individual ALON001 carrying haplogroup E1b1b1a1, which likely originated in North Africa before spreading to the Mediterranean area of Europe (Battaglia *et al*., 2009). This finding indicates a male contribution to the previously identified European ancestry in the genome of this individual.

The prevalence and frequency of mtDNA haplogroups differ significantly between regions and times. In Argentina, mtDNA haplogroup frequencies follow a spatial pattern. In Northwest Argentina and Northwest Patagonia, haplogroups D1g and C1b13 are reported at the highest frequencies in present-day populations (Postillone *et al*., 2020b). These two clades are identified at lower frequency in our time transect from the region. Instead, ancient individuals from the Pampas region show the co-occurrence of D1g and D1j haplogroups, besides the presence of the major A2, B2, C1 lineages and their sub-lineages (Postillone *et al*., 2020b; Roca-Rada *et al*., 2021; Motti *et al*., 2023). These two lineages show rare occurrences in present-day Chile and Argentina, but local high frequencies (Bodner *et al*., 2012). We detected four Pampas individuals in our dataset with D1g, indicating its persistence in the region through time (Table S1).

For Uruguay, published ancient mtDNAs reveal three different haplogroups, namely C1d3, C1b, and C1c (Sans *et al*., 2015; Figueiro *et al*., 2017; Lindo *et al*., 2022), while modern-day Uruguayans revealed the prevalence of both B2 and C1 haplogroups (Spangenberg *et al*., 2021). Consistent with these results, the ancient individuals from the eastern lowland of Uruguay (INDI_870BP) and from the Lower Uruguay River (LOCA_1600BP) carry mtDNA haplogroups C1b and B2, respectively.

Genetic kinship within the new genomic dataset was estimated using KIN (Popli *et al*., 2023). This revealed TPAG001 and TPAG004, buried in the same mound in the Paraná River Delta, as parent-child related, and multiple second- and third-degree relationships were identified both within and across sites in the southern Pampas (eastern Pampa-Patagonia transition) and Northwest Patagonia (Table S1).

We calculated the runs of homozygosity (ROHs) for all individuals with sufficient data (Material and Methods). We found no evidence of routinely practiced close-kin mating patterns at any of the sites. However, an increased number of long ROHs (20-300 cM) in LATC_3100BP from the central Pampas and AQUI005_4200BP from Northwest Patagonia indicates that their parents were closely related (Figure S2). Our results suggest evidence of generally low effective population sizes as revealed by the high number of short ROHs (4-8 cM), which tend to decrease over time. This fits with the archaeological evidence for the Pampas, where smaller hunter-gatherer groups persisted until the Late Holocene, when larger populations appeared (Politis & Borrero, 2024).

### Population genetic analysis overview

For population genetic analyses, we grouped the individuals based on the archaeological site, date, and statistical fit within the tested groups; otherwise, we analysed them separately (Tables S1 and S2). We removed individuals involved in first- and second-degree relationships. Finally, we combined the newly generated data with ancient (1240K dataset) and modern-day (Illumina masked dataset) Indigenous American genome-wide data (Reich *et al*., 2012; Mallick *et al*., 2016; Mallick *et al*., 2024) (Materials and Methods). To gain an overview of the genetic relationships between the newly sequenced individuals and other Southern Cone populations, we built a multidimensional scaling (MDS) plot based on 1-f_3_-outgroup statistics (Figure 2A). By plotting the first two components, we obtained a MDS plot that broadly mirrors geography, with most central and southern Pampas individuals clustering on the bottom right, most Northwestern Patagonia individuals on the bottom left, close to previously published southern Andean individuals (LosRieles_5100BP and Conchali_700BP), and all Paraná River Delta and Uruguay individuals on top. Most of the oldest groups from both the central Pampas (ArroyoSecoII_7700 BP and LagunaChica_6800BP) and Northwestern Patagonia (AQUI_4200BP) fall in a central position of the plot. Interestingly, five groups from Northwestern Patagonia (buried in the east of Neuquén) fall in proximity to the primarily Pampas cluster, suggesting at least two distinct ancestries present in this region through time. One Middle Holocene group from the southern Pampas at the eastern Pampa-Patagonia transition (TREB_5200BP) is shifted towards the Northwest Patagonia/southern Andean group. However, an extended MDS plot reveals not only TREB_5200BP, but also REMO_230BP, to be shifted towards ancient southern Patagonian populations (Figure S3).

**Figure 2:**
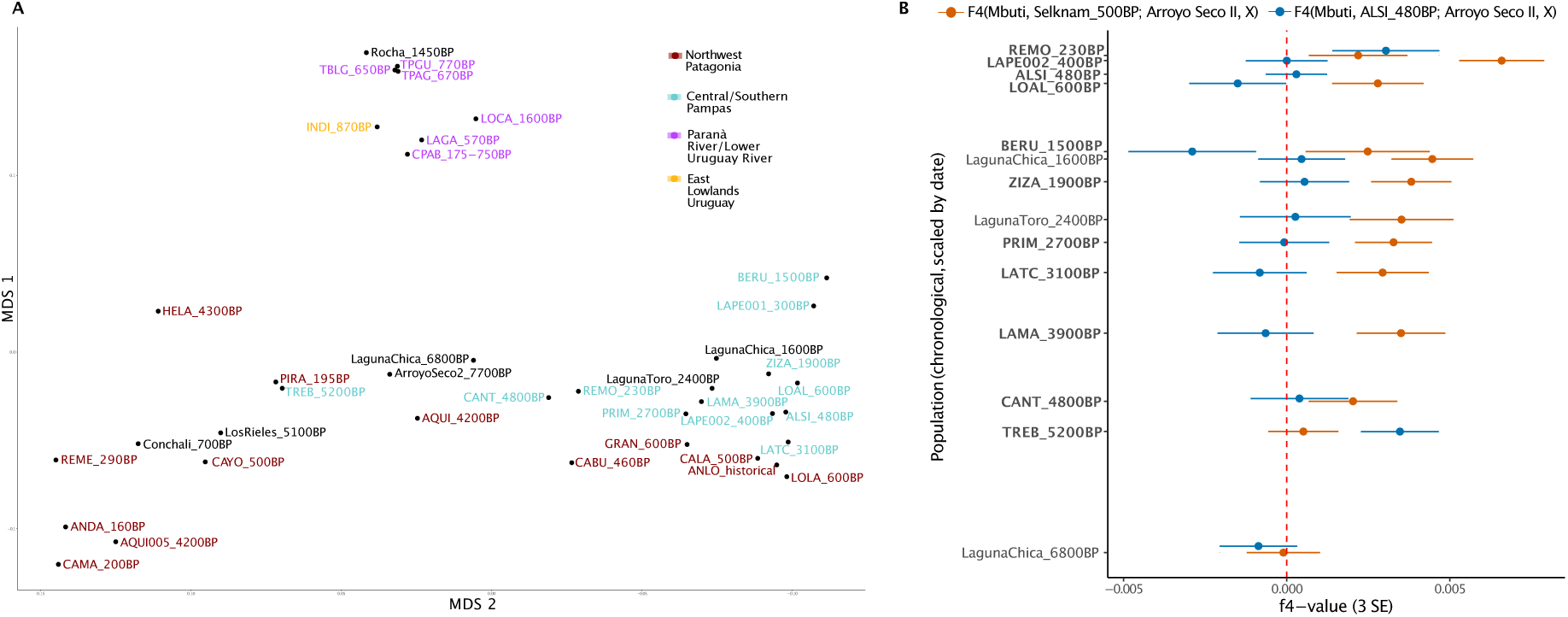
Genetic similarity of ancient southern Pampas individuals/ groups to modern-day and ancient Indigenous Americans. A) Multidimensional scaling plot based on 1-f3 outgroup statistics on the 1240K panel including our newly generated data and representative published populations from the southern Andes, the Pampas, and Uruguay. B) f4-statistics plot. The y-axis has been sorted chronologically proportional to the date of the individuals/populations (newly reported in bold). Each population has been tested for specific affinity to southern Andean ancestry represented by Conchali_700BP (blue), the ancestry of unknown geographic origin represented by ALSI_480BP (orange), and southern Patagonian ancestry represented by Selknam_500BP (blue), against an ancestry baseline for the Pampas represented by ArroyoSecoII_7700BP. f4-values are depicted with 3 standard errors (SE).

### At least three distinct ancestries in Middle Holocene Pampas

Our dataset comprises genomic data of 21 individuals from eleven sites in central and southern Pampas spanning about 5,000 years, with the oldest individuals dated to 5,200 BP and the youngest to 200 BP (Table S1 and Supplementary Information).

We first tested for the general affinity of newly and previously reported Pampas genomes with modern-day Indigenous American populations (X) through f_3_-outgroup statistics of the form f_3_(Mbuti; X, Pampas) (Table S3.1). A previous study has established a significant affinity of the oldest central Pampas individuals (ArroyoSecoII_7700BP and LagunaChica_6800BP) with modern-day populations from the southern region of the Southern Cone (Posth *et al*., 2018). Interestingly, TREB_5200BP, REMO_230BP, and, to some extent, also CANT_4800BP, at the eastern Pampa-Patagonia transition (southern Pampas), show a similar pattern of genetic affinity to multiple present-day populations from southern Chile and the southern tip of Patagonia (Figure S4). Instead, all other Pampas populations exhibit a more even distribution of genetic similarity to modern-day populations from northern, central, and southern South America, indicating that they primarily derive from a distinct ancestry not fully represented in the modern-day populations available in our dataset.

We then compared previously and newly sequenced ancient individuals from the Pampas to ancient Indigenous American populations (X) with f_3_(Mbuti; X, Pampas) (Table S3.2). Most newly reported central and southern Pampas individuals show more allele sharing with 4,800-230 BP groups from the same areas, including previously published Late Holocene genomes such as LagunaToro_2400BP and LagunaChica_1600 BP (Nakatsuka *et al*., 2020a; Nakatsuka *et al*., 2020b). Instead, the oldest and youngest individuals in the southern Pampas time transect (TREB_5200BP and REMO_230BP) exhibit the highest genetic affinity to ancient populations from southern Patagonia.

We then investigated how our newly generated genomes from the central and southern Pampas relate to previously sequenced genomes from these areas by conducting f_4_-statistics of the form f_4_(Mbuti, X; Pampas 1, Pampas 2) (Table S3.3). When X is ArroyoSecoII_7700BP or LagunaChica_6800BP, comparisons generally show Z=0, indicating equal levels of allele sharing between younger central and southern Pampas individuals in comparison to the oldest Pampas genomes. Instead, when X is LagunaChica_1600BP or LagunaToro_2400BP, f_4_-statistics result in Z>|3| for individuals dated between 3900 and 400 BP against TREB_5200BP and REMO_230BP, indicating excess allele sharing between Late Holocene central and southern Pampas individuals, to the exclusion of the oldest and youngest individuals in our southern Pampas dataset. Thus, the ancestry previously reported for LagunaChica_1600BP and LagunaToro_2400BP is not restricted to these individuals but widespread in the central and southern Pampas at least by the beginning of the Late Holocene.

We used ALSI_480BP as a representative of this ancestry of unknown geographic origin since it is the population from the southern Pampas with the highest number of individuals (n=9) that maximises the signal in the previous f_4_-statistics. We then implemented another set of f_4_-statistics to more precisely detect the arrival of the incoming ancestry in relation to the baseline genetic profile of ArroyoSecoII_7700BP, with the following configuration f_4_(Mbuti, ALSI_480BP; ArroyoSecoII_7700BP, Pampas) (Figure 2B, Table S3.4). These statistics show a significantly higher (Z>|3|) allele sharing between ALSI_480BP and all ancient Pampas individuals/groups compared to ArroyoSecoII_7700BP. Only the second two oldest individuals from the region, i.e. LagunaChica_6800BP and TREB_5200BP, result in statistics consistent with 0. Interestingly, CANT_4800BP and REMO_230BP also provide significant statistics, indicating that the incoming ancestry arrived in the region before the end of the Middle Holocene and survived until colonial times. Nevertheless, we observe that the contribution of this ancestry was not constant through time. With a series of f_4_-statistics, we reveal an additional increase in the incoming ancestry by 3,900 BP, and possibly another surge by 400 BP, suggesting a prolonged period of gene-flow from an ancestry of unknown geographic origins into the central and southern Pampas (Table S3.4).

The lack of ALSI_480BP-related signal in the three oldest Pampas groups (ArroyoSecoII_7700BP, LagunaChica_6800BP, TREB_5200BP) when compared against one another (Figure 2B, Tables S3.4, 3.5) alongside their similar affinity to present-day populations shown in f_3_-outgroup statistics (Figure 2A) raise the possibility that these Middle Holocene populations share the same ancestry and derive from a genetically homogeneous group. We specifically tested for population continuity between ArroyoSecoII_7700BP, LagunaChica_6800BP and TREB_5200BP by conducting a series of f_4_-statistics of the form f4(Mbuti, X; oldest Pampas, oldest Pampas) where X are ancient Indigenous American genomes. When directly comparing ArroyoSecoII_7700BP to TREB_5200BP, the latter shows significant affinity to ancient southern Patagonian populations from Chile and Argentina. Instead, when directly comparing LagunaChica_6800BP and TREB_5200BP, the former shares more drift with CANT_4800BP. However, LagunaChica_6800BP and CANT_4800BP are not cladal groups since a direct comparison shows a significantly higher affinity of CANT_4800BP to the incoming ancestry represented by ALSI_480BP (Table S3.6). Taken together, these results reject the hypothesis of complete genetic continuity among the available Middle Holocene genomes from the central and southern Pampas but indicate the presence of at least three genetically distinct ancestries related to 7,700-6,800 BP central Pampas, southern Patagonia by 5,200 BP, southern Pampas by 4,800 BP, the latter becoming the dominant ancestry in both the central and southern Pampas from 3,900 BP onwards.

To investigate if the southern Patagonian gene flow is observed in the central and southern Pampas through time, we extended the previously tested configuration of f_4_-statistics to f_4_(Mbuti, Selknam_500BP; ArroyoSecoII_7700BP, Pampas) (Figure 2B). As expected from the conducted f_3_-outgroup statistics, we identify a significant affinity to southern Patagonian ancestry both in TREB_5200BP and REMO_230BP. The detection of this signal in the oldest and youngest individuals in our newly generated dataset suggests an intermittent presence of southern Patagonian ancestry into the southern Pampas (eastern Pampa-Patagonia transition). Notably, two Pampas individuals in the dataset, i.e., BERU_1500BP and LOAL_600BP, show a significantly lower affinity to the southern Patagonian ancestry in comparison to ArroyoSecoII_7700BP (Figure 2B). This might suggest their admixture with yet other ancestral sources in the genetic history of these two individuals or undetected levels of present-day contamination.

A previous study had revealed a clear genetic distinction between ancient populations from southern Patagonia that mirrors their geography distribution and subsistence strategy (Nakatsuka *et al*., 2020b). Specifically, ancient populations in southwestern Patagonia (Western Archipelago and Beagle Channel) who primarily relied on marine resources were found to be genetically distinct from ancient populations in southeastern Patagonia (Mitre Peninsula, Northern Tierra del Fuego and Southern Continent) who based their diet primarily on terrestrial resources. To better understand the relationship between TREB_5200BP, REMO_230BP and populations from southern Patagonia, we constructed a set of comparative f_4_-statistics of the form f_4_(Mbuti, TREB_5200BP/REMO_230BP; southern Patagonia 1, southern Patagonia 2). These revealed a clear affinity of TREB_5200BP and REMO_230BP to Late Holocene populations from southeastern rather than southwestern Patagonia (Table S3.7). We were finally able to model TREB_5200BP in qpAdm as a mixture between 37±13.5% ArroyoSecoII_7700BP and 63±13.5% Selknam_500BP, as proxies for the local and incoming southeastern Patagonia ancestries, respectively (Table S4). These results suggest either long-distance migrations from southeastern Patagonia into the southern Pampas or, alternatively, that this type of ancestry was more widespread and in geographic proximity to the Pampa-Patagonia transition.

### Persistence of Andean ancestry in Northwest Patagonia

We report a separate time transect of 4,300 years in Northwest Patagonia, from the Neuquén province of Argentina, closely located to the Chilean border and the Andean Mountain range. The previously presented MDS plot (Figure 2A), showed the distribution of Northwest Patagonian populations into two clusters. Specifically, seven groups, mainly from Northwest Neuquen and close to Andean mountain, cluster in proximity to southern Andean individuals (HELA_4300BP, AQUI005_4200BP, CAYO_500BP, REME_290BP, CAMA_200BP, PIRA_195BP, ANDA_160BP), while the other five cases, mainly located close to the Limay River in eastern Neuquén, cluster close to 4,800-230 BP Pampas individuals (LOLA_600BP, GRAN_600BP, CALA_500BP, CABU_460BP, ALON_historical). Interestingly, one of the oldest groups, AQUI_4200BP, falls in an intermediate position along these two clusters. This pattern suggests the presence of multiple ancestries in Northwest Patagonia from the end of the Middle Holocene to the end of the Late Holocene.

As previously done for the Pampas time-transect, we first conducted f_3_-outgroup statistics of the form f_3_(Mbuti; X, Northwest Patagonia) (Table S5.1), where X are modern-day Indigenous American populations (Figure 3A, Figure S5). Northwest Patagonian groups that cluster in the MDS plot close to ancient southern Andean individuals, show excess affinity to modern-day central and southern Patagonian populations such as Hulliche, Chilote, Yaghan and Chono (Figure 3A). Instead, groups that fall close to 4800-230 BP central and southern Pampas individuals, show a more even distribution in their affinity to all central and southern Indigenous Americans in our dataset (Figure 3A).

**Figure 3:**
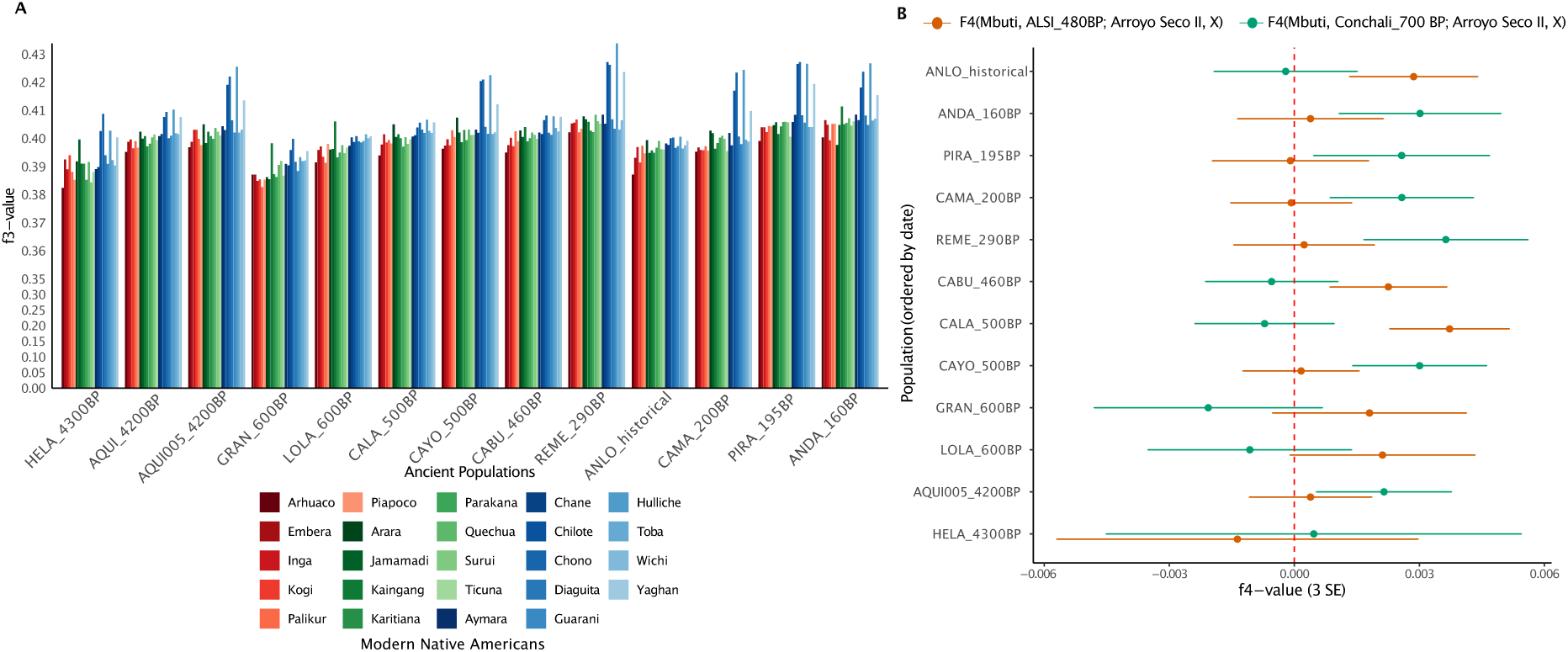
Genetic similarity of Northwest Patagonia populations to present-day and ancient Indigenous Americans. A) Bar-plot based on f3-outgroup statistics of the form f3(Mbuti; Northwest Patagonia, X), where X are modern-day populations that have been genotyped on the Illumina panel in which European and/or African ancestries have been masked (Reich *et al*., 2012). Representatives of each major South American region have been selected, sorted and color-coded following a north-to-south order. The X-axis represents the chronological time transect of the Northwest Patagonia region, with newly reported individuals/groups in bold font. The Y-axis has been collapsed between 0 and 0.35 and extended between 0.35 and 0.43 for better resolution. B) f4-statistics plot. The y-axis has been sorted chronologically from older to younger individuals/populations (not in scale). Each population has been tested for specific affinity to southern Andean ancestry represented by Conchali_700BP (green) and the ancestry of unknown geographic origin represented by ALSI_480BP (orange) against the ancestry baseline for Northwest Patagonia represented by AQUI_4200BP. f4-values are depicted with 3 standard errors (SE).

To formally test the clustering pattern observed in f_3_-outgroup statistics we conducted a series of f_4_-statistics of the form f_4_(Mbuti, X; AQUI_4200BP, Northwest Patagonia) (Table S5.2), assuming AQUI_4200BP to represent the baseline ancestry of the region, while X being ALSI_480BP or Chile Conchali_700BP to represent the ancestry of unknown origins that reached the Pampas by 4800 BP or the southern Andean ancestry, respectively (Figure 3B). In the comparisons with ALSI_480BP, Z exceeds the significance threshold for all groups falling close to 4800-230 BP central and southern Pampas individuals in the MDS plot (Figure 3B, Figure 2A). Instead, when compared to Conchali_700BP, f_4_-statistics are significant for all groups previously shown to cluster near ancient southern Andean individuals (Figure 2A) and with the highest f_3_-outgroup statistics values for modern-day southern Patagonian populations (Figure 3A). These results are confirmed with f_4_(Mbuti, Northwest Patagonia; Conchali_700BP, ALSI_480BP). The Northwest Patagonian individuals previously identified to carry the ancestry of unknown origins are significantly closer to ALSI_480BP, while the ones with southern Andean ancestry are significantly closer to Conchali_700BP (Table S5.3).

We finally investigated possible genetic links between late Middle Holocene and Late Holocene individuals in Northwest Patagonia with f_4_(Mbuti, Middle Holocene; Late Holocene 1, Late Holocene 2) (Table S5.4). The results for AQUI_4200BP and HELA_4300BP do not produce significant values deviating from zero, suggesting that all Late Holocene individuals are equally related to them. However, the comparisons with AQUI005_4200BP show higher allele sharing with CAMA_200BP, ANDA_160BP and REME_290BP. This suggests that the southern Andean ancestry present at the end of the Middle Holocene (AQUI005_4200BP) in Northwest Patagonia persisted until the end of the Late Holocene.

In summary, our results reveal that the Middle and Late Holocene population history of Northwest Patagonia involved at least three genetic lineages: the first related to 4,300-4,200 BP Northwest Patagonia, the second related to the southern Andes as early as 4,200 BP, and the third one by 600 BP representing the same unknown ancestry that reached the southern Pampas by 4,800 BP.

### Local structure along the Paraná River Delta and Uruguay coastal connections

Our newly generated dataset also encompasses five Late Holocene sites from the Paraná River Delta in Argentina, two from the Upper Delta of the Paraná River (CPAB and LAGA) and three from the Lower Delta of the Paraná River (TPAG, TPGU, TBLG), dated between 770 and 175 BP. Additionally, we obtained genomic data from two ancient Uruguayan individuals, one from the Lower Uruguay River dated to 1,600 BP (LOCA), and the other from the eastern lowlands of Uruguay dated to 870 BP (INDI).

We conducted two sets of f_3_-outgroup statistics of the form f_3_(Mbuti; Paraná River/Uruguay, Modern Indigenous Americans) and f_3_(Mbuti; Paraná River/Uruguay, Ancient Indigenous Americans) to understand potential affinities of these individuals/groups to ancient and present-day Indigenous American populations. The first test revealed excess allele sharing between all tested populations and the present-day Kaingang, a Je-speaking population from Brazil (Figure S6). We formally tested this genetic connection with f_4_(Mbuti, Paraná River/Uruguay; Kaingang, Modern South Americans). However, to the limit of our resolution, we were unable to establish any significant affinity of individuals from Uruguay and the Paraná River to present-day Kaingang (Tables S6.1 and 6.2).

In the second set of f_3_-outgroup-statistics, individuals from the Upper and Lower Delta of the Paraná River show the highest affinity among themselves. More specifically, individuals from different Upper Delta sites tend to be genetically closer to other Upper Delta individuals, and the same is true for individuals from Lower Delta sites. These similarities create a clustering based on geography that is also visible in the MDS plot (Figure 2A). Additionally, groups from the Upper and Lower Delta of the Paraná River, and also LOCA_1600BP from Lower Uruguay River, exhibit a general similarity to 4800-230 BP central and southern Pampas individuals, who were described to carry an ancestry of unknown geographic origin. We confirmed the presence of this ancestry also in the Paraná River Delta with the f_4_-statistics f_4_(Mbuti, Paraná River/LOCA_1600BP; ArroyoSecoII_7700BP, ALSI_480BP) that provided Z-scores >|3| for all but one test (Table S6.3).

In f_3_-outgroup statistics we observe high allele sharing of Paraná River Lower Delta and Uruguayan individuals with post-2,200 BP Sambaqui-associated groups, previously shown to be related to the spread of Kaingang ancestry to the southern coast of Brazil (Ferraz *et al*., 2023). We then tested for this specific affinity with f_4_(Mbuti, Paraná River/Uruguay; ArroyoSecoII_7700BP, Kaingang 100BP) (Table S6.4). All Z-scores are consistent with 0 for Paraná River groups and LOCA_1600BP. Instead, we obtain a Z-score >|3| for INDI_870BP, indicating genetic links between the coastal expansions to southern Brazil and Uruguay.

We finally constructed a neighbour-joining tree based on inverted f_3_-outgroup statistics including representative genomes from the central and southern Pampas, Northwest Patagonia, southern Andes and southern coast of Brazil, as well as all newly generated individuals/groups from the Paraná River Delta and Lower Uruguay River, and eastern lowlands of Uruguay (Figure 4). The tree topology illustrates the previously described geographic clustering with the Lower Paraná River populations forming a clade. Importantly, LOCA_1600BP clusters as a sister group to the Lower Paraná River populations, indicating a level of genetic continuity in the region for more than a millennium. Instead, both INDI_870BP and the previously published Rocha_1450BP, both from the Eastern lowlands of Uruguay, cluster together on a branch with Sambaqui-associated individuals from southern Brazil.

**Figure 4:**
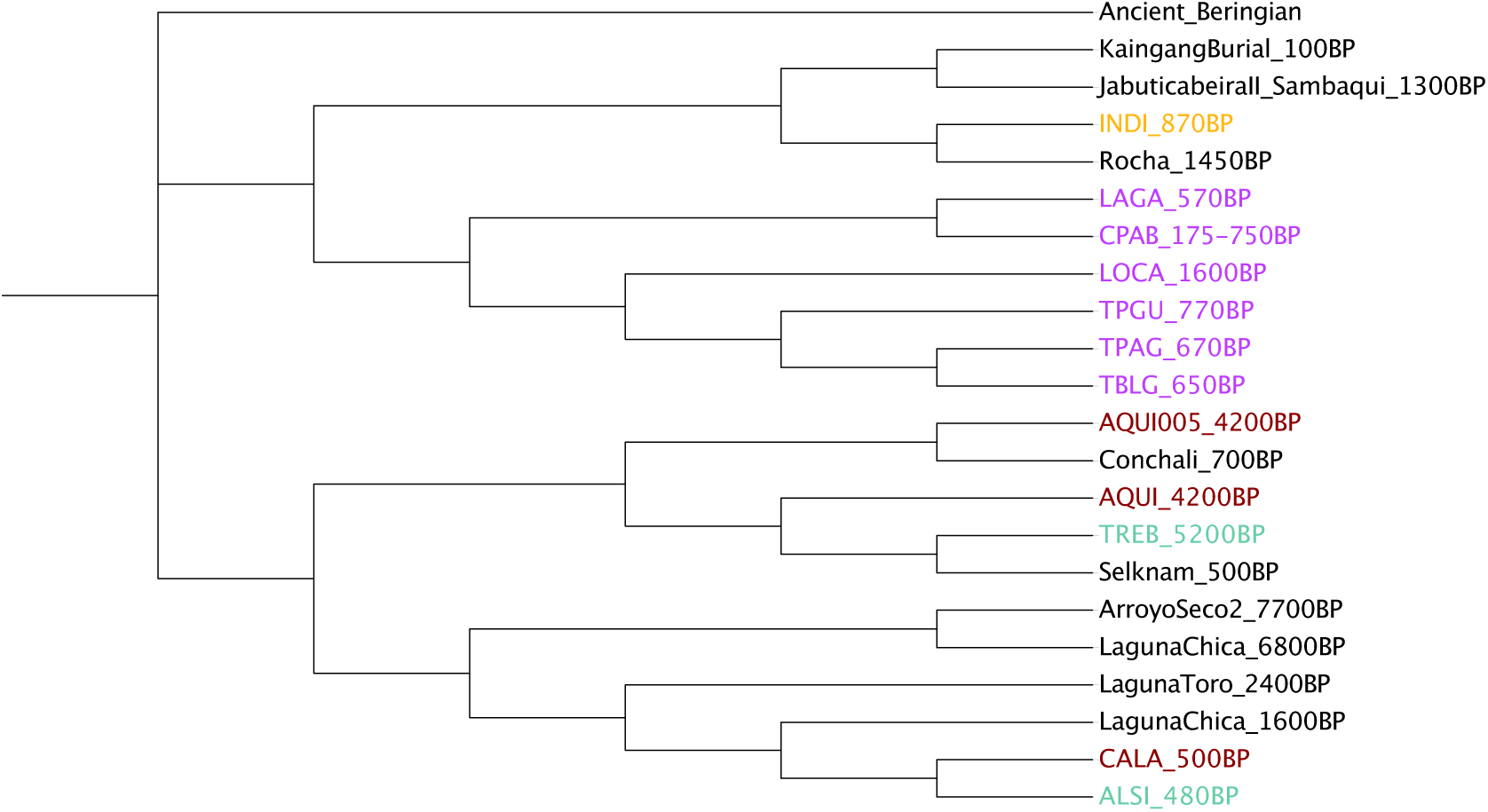
Neighbour-joining tree showing the genetic clustering of ancient CSC populations. The tree is built with inverted f3-values (1/f3). All individuals/groups from the Paraná River Delta and Lower Uruguay River and from the eastern lowlands of Uruguay are represented. Instead, a subset of individuals/groups from Northwest Patagonia and central and southern Pampas have been selected as representative of the main ancestries present in those regions during the Middle and Late Holocene. Southern Brazilian, southern Patagonian, and southern Andean ancient individuals have been selected based on significant affinities with CSC populations obtained in the previously described f4-statistics. Branch lengths have been disabled and label colours for newly reported individuals are based on geography, as in all previous figures.

## Discussion

In this study, we present mtDNA and genome-wide data from 52 individuals across four geographical regions in SCS (central and southern Pampas, Northwest Patagonia, Paraná and Uruguay Rivers, and eastern Uruguay), spanning a time transect of about five millennia, and filling important temporal and geographic gaps in the archaeogenomic record of the Southern Cone of South America. Our findings reveal a complex population history, with genetic connections between distinct ancestries across all studied regions.

### Central and Southern Pampas

It was suggested that the ancestry found in ArroyoSecoII_7700BP, and LagunaChica_6800BP, was characteristic of the Pampas during the Middle Holocene (Posth *et al*., 2018). Our newly reported data, however, suggest that this is only one of at least three distinct ancestries present in the Middle Holocene Pampas. We showed that two individuals from the Tres Bonetes site (TREB_5200BP) in the southern Pampas (eastern Pampa-Patagonia transition) exhibit an excess affinity to southern Patagonian ancestry. More specifically, this genetic component is distinctive for populations from the eastern region of southern Patagonia. Thus, we modelled the Tres Bonetes individuals as resulting from an admixture between local (ArroyoSecoII_7700BP) and incoming (Selknam_500BP) ancestries. Due to the lack of ancient genomes from central Patagonia, we are unable to assess if the ancestry identified in Tres Bonetes is the result of a long-distance south-to-north migration from southeastern Patagonia, or if it was also distributed more closely to Pampas. Interestingly, we find a similar genetic connection with southeastern Patagonian populations in the most recent individual of our southern Pampas time transect (REMO_230BP).

We proposed two possible explanations for the non-continuous presence of the southeastern Patagonian ancestry in the southern Pampas: 1) it co-existed with other lineages but remains largely unsampled; 2) it was the result of independent gene flows into the region. At least for historical periods, there are evidence of south-to-north migrations from Gunnuka-kena or Aonikenk (also known as the northern and southern Tehuelche ethnolinguistic groups, respectively) into the Pampas region (Casamiquela, 1965; Nacuzzi, 1998; Alioto, 2011; Roulet, 2016). Thus, the southeastern Patagonian-like ancestry might have been present in or in the vicinity of the Pampas, rather than arriving with long-range migrations from the southern tip of South America.

Lastly, we describe a third ancestry in the Middle Holocene southern Pampas. To the limit of our resolution, constrained by available ancient genomes from South America, we are unable to identify a proxy population for this ancestry outside the Pampas before the Late Holocene, making it currently impossible to infer its geographic origin. To indirectly trace the arrival and temporal distribution of this genetic component in the central and southern Pampas, we used genomes from the Paso Alsina group (ALSI_480BP). Through f-statistics and MDS analyses, we not only showed the arrival of this ancestry in the southern Pampas by 4,800 BP, but also observed a significant increase in this component by 3,900 BP. These results provide strong support for a change in the genetic composition of the central and southern Pampas populations due to a gene flow starting in the Middle Holocene from a genetically distinct population. Thus, we do not find evidence for a complete population replacement as previously proposed for the southeastern Pampas between the Middle and Late Holocene (Barrientos & Pérez, 2004; Barrientos & Perez, 2005; Barrientos, 2009). This period is followed by a phase of broad genetic continuity without significant ancestry influx at least until 400 BP, which might be consistent with a model of population and cultural continuity during the Late Holocene (Martínez *et al*., 2024; Politis & Borrero, 2024).

### Northwest Patagonia

The newly generated data from Northwest Patagonia derive from the Neuquén province of Argentina, located along the southern Andes. Our results support a significant southern Andean genetic influence in Northwest Patagonia from around 4,300 BP onwards as baseline ancestry (HELA_4300BP and AQUI_4200BP). Interestingly, another similarly dated individual from the Aquihueco site (AQUI005_4200BP) shows an even higher affinity to the southern Andes, suggesting this individual as a recent migrant into the region.

Since our time transect in Northwest Patagonia has a large gap until the terminal phase of the Late Holocene (Figure 1B), we cannot reject hypotheses of population continuity or discontinuity from the Middle to Late Holocene. Nevertheless, post-600BP individuals from east Neuquén tend to cluster with 4800-230 BP central and southern Pampas individuals and exhibit a higher affinity to ALSI_480BP than AQUI_4200BP. This indicates the ancestry of unknown geographic origin, which arrived in the Pampas by 4,800 BP, also reached Northwest Patagonia sometime between 4,200 BP and 600 BP. It is important to note that this gene flow was not identified in Andean individuals from western Neuquén but in individuals from eastern Neuquén located along the Negro and Limay Rivers, which flow to the Atlantic coast. Thus, an alternative and perhaps more likely explanation for the attraction of post-600BP Northwest Patagonia individuals to the post-4,800 BP central and southern Pampas individuals is a migration from the Pampas directly into Northwest Patagonia. Finally, the extra southern Andean affinity identified in AQUI005_4200BP is found again from 500 BP in the Neuquén Andes and seems to co-exist side-by-side with the Pampas-like ancestry until post-colonial times.

The obtained genetic results have possible matches with models for the population history of Northwest Patagonia put forward through the study of the archaeological record. The arrival of allochthonous ancestries in the region as late as 600 BP might be related to archaeological transitions observed during the final phase of the Late Holocene, when forager lifestyles gradually were replaced by more sedentary forms of residence. Similarly, the presence of both the Pampas- and southern Andean-related ancestries around the same time might be associated with the increase in differentiation of regional cultural developments during the Late Holocene (Perez *et al*., 2016; Cobos *et al*., 2022).

### Paraná/Uruguay Rivers and eastern lowlands of Uruguay

The Paraná River Delta and the Lower Uruguay River are located between the Pampas, Uruguay and southern Brazil, which are regions with newly generated and previously published ancient genomic data (Moreno-Mayar *et al*., 2018; Posth *et al*., 2018; Ferraz *et al*., 2023). This allowed us to investigate whether these two regions were within the genetic spheres of neighbouring populations or if they carried a yet distinct ancestry. We report 15 individuals from six sites spanning 1,600 years in the Paraná River Delta and Lower Uruguay River, as well as one individual from the eastern lowlands of Uruguay dated to 870 BP. Testing for affinity to modern-day Indigenous American populations revealed no consistent genetic attraction toward any population available in our dataset.

However, when comparing the newly generated genomic data against ancient Indigenous American populations, individuals from the Upper and Lower Delta of the Paraná River, as well as a genome from the Los Cardos site in the Lower Uruguay River (LOCA_1600BP) exhibit a distinct affinity to post-4,800 BP Pampas individuals as well as to individuals from Northwest Patagonia clustering with the same Pampas groups in the MDS plot. Therefore, the migration that brought the ancestry of unknown geographic origins into the Pampas and Northwest Patagonia between 4,800 BP and 600 BP, respectively, also reached the Lower Uruguay River and Paraná River Delta by 1,600 BP.

When conducting f_3_-outgroup statistics comparing Paraná River Delta populations with modern and ancient Indigenous Americans, other Paraná River Delta populations were always the best match, suggesting a rather homogeneous genetic makeup. This homogeneity, suggestive of a shared ancestry, is mirrored in a common settlement pattern involving earthen mound constructions and in a broadly similar subsistence economy centered on aquatic faunal resources and supplemented by small-scale horticulture.

However, in MDS space (Figure 2A), ancient populations from the Paraná River Delta fall into two separate clusters that do not follow a temporal pattern but rather a geographic one (Upper and Lower Delta). These genetic differences are paralleled by variations in ceramic technological styles between the two sub-regions, indicating that they represent separate archaeological entities (Serrano, 1950; Ceruti, 2003; Politis & Bonomo, 2023). Most interestingly, this genetic structure appears to result from differential influxes of ancestry into the Lower and Upper Delta sections. f_4_-statistics reveal that Lower Paraná River Delta individuals harbour an extra affinity to INDI_870BP, an individual from the eastern lowlands of Uruguay. In contrast, Upper Paraná River Delta individuals share more genetic drift with LOCA_1600BP, which is geographically much closer to the Lower Delta of the Paraná River. This genetic observation aligns with cultural evidence indicating the Goya Malabrigo archaeological entity, shared between the Upper Paraná River Delta and the Lower Uruguay River, as evidenced at the Los Cardos site.

When comparing both Uruguayan individuals to a previously published 1450-year-old genome from the Rocha site in eastern Uruguay (Lindo *et al*., 2022), we find that INDI_870BP is significantly closer to the Rocha individual than LOCA_1600BP. This confirms a clear geographic pattern where populations from the eastern lowlands of Uruguay are more closely related to each other than to a genome from inland Uruguay.

Importantly, INDI_870BP consistently shows attraction to post-2,200 BP individuals from southern Brazil associated with the Sambaqui archaeological culture. An earlier study showed that these individuals carry Kaingang-related ancestry, which has been linked to the expansion of Jê languages (Ferraz *et al*., 2023) and to extensive earthwork constructions (Iriarte *et al*., 2017). Our newly sequenced individual (INDI_870BP) is, so far, the only one from Uruguay found in association with the mound builder archaeological tradition, which shows archaeological connections with tropical lowland and coastal cultures (Andrade Lima & López Mazz, 1999). The match between the archaeological and genetic findings suggests that the spread of Kaingang-associated ancestry continued southward along the coast of southern Brazil, reaching Uruguay by at least 870 BP. Interestingly, Uruguayan modern-day individuals with high levels of Indigenous American ancestry also show a genomic affinity with Kaingang groups (Spangenberg *et al*., 2021). However, due to our limited and sparse sampling, further sampling efforts are needed to confirm the differential genetic patterns observed in this study across different regions of present-day Uruguay.

## Conclusions

In conclusion, we present three key insights. First, we identify the expansion of an ancestry of unknown geographic origins into three CSC regions–appearing by 4,800 BP in the Pampas, by 1,600 BP in the Lower Uruguay River and by 600 BP in Northwest Patagonia– events that reshaped the genomic landscape of these areas. Second, we detect repeated population movements into the southern Pampas and Northwest Patagonia involving individuals with ancestries related to the southern Andes and southeastern Patagonia, respectively. These findings indicate a high degree of interconnectivity among distant regions of the Southern Cone in the Middle and Late Holocene, until colonial times. Third, we uncover close genetic affinities between the eastern lowlands of Uruguay and Sambaqui groups, suggesting the coastal dispersal of an ancestry related to Jê language speakers from southern Brazil to eastern Uruguay in association with mound builder societies.

Thus, our study reveals that large-scale human movements across the Southern Cone of South America were closely linked to major cultural changes during the Middle and Late Holocene. While our dataset is temporally and geographically vast and triples the number of ancient individuals with reported genome-wide data from the region, additional archaeogenomic studies are necessary to build a more holistic understanding of the historical dynamics of Indigenous populations in the Southern Cone of South America.

## Material and Methods

### Archaeological sampling

Permits to perform genetic research on ancient individuals from Argentina and Uruguay were granted by the Dirección de Cultura (Entre Ríos), Ministerio de Innovación y Cultura (Santa Fe), Museo de La Plata, Ministerio de las Culturas, Gobierno de la provincia del Neuquén, Dirección Provincial de Patrimonio Cultural del Gobierno de la Provincia de Buenos Aires and the Instituto Nacional de Pensamiento Latinoamericano (INAPL), Comisión del Patrimonio Cultural de la Nación and the Museo Nacional de Antropología de Montevideo.

Overall, we obtained genomic data from 52 individuals in dedicated ancient DNA laboratories at the University of Tübingen (n=33), University of Pavia (n=14), and Harvard Medical School (n=5). A detailed description of the samples processed in each laboratory is provided in Table S1.

### DNA extraction

At the University of Tübingen, each petrous portion of the temporal bone and each tooth was photographed from multiple angles before destructive sampling. We cut teeth at the dentine-enamel junction using an electric saw blade. We generated dentine powder by sampling the crown pulp with an electric dental drill. Petrous bones were sampled by abrasing the outermost layer of the bone surface before sampling the cochlea from the internal acoustic meatus. We produced 30-64 mg of bone/tooth powder that was used for DNA extraction. To digest the bone matrix, 1 ml of a buffer containing 0,45M of EDTA [0.5 M, pH 8], UV-water, as well as 0,25 mg/ml Proteinase K, was added to each sample. This was followed by an overnight incubation step at 37°C with constant rotation. The undigested remains of bone powder were pelleted, and the supernatant was transferred into a tube containing 10 ml of UV-treated binding buffer [GuHCL 5M, Isopropanol 40%, UV-treated H_2_O] and 400 µl Sodium acetate [3M, pH 5.2]. Each sample was then transferred into a 50 ml Falcon tube containing a silica column for high-volume purification (High Pure Viral Nucleic Acid Large Volume Kit; Roche) (Dabney *et al*., 2013; Korlević *et al*., 2015). The purified DNA was eluted in 2x 50 µl tris-EDTA-Tween (TE buffer, 0.05% Tween) and stored in a dedicated freezer at - 20°C until further use At the University of Pavia, a photograph of each sample was taken before destructive sampling. Each petrous portion of the temporal bone was cut in half to remove the spongy bone surrounding the cochlea. Afterwards, the cochlea was perpendicularly cut, and half of it was pulverised. Bone samples were pulverised in a Retsch MM400 mixer mill in zirconium oxide grinding jars and between 50 and 100 mg bone powder was obtained and was used for ancient DNA extraction. For DNA extraction, we followed a modified version of a published protocol (Rohland *et al*., 2018). Specifically, 1 ml lysis buffer containing 1 ml EDTA [0.5 M, pH 8], 20 mg/ml Proteinase K, and 10% (mg/ml) N-lauroylsarcosine was added before an overnight incubation at 50°C with constant rotation. Undigested remains of bone were pelleted, 1 ml of the supernatant was transferred into 5 ml of binding buffer [Quiagen] and incubated at room temperature for five minutes. The mixture is then transferred into a 50 ml Falcon tube containing a silica column for high-volume purification (Minelute). The purified DNA is eluted in prewarmed EBT buffer (10 mM Tris-HCl, pH 8.5, 0,05% Tween) first for 55 µl, then 2x 22 µl and stored at -20°C.

At Harvard Medical School, around 50 mg of bone and tooth powder was generated using an electric dentist drill. DNA was extracted from powdered samples using a method optimized for retaining small DNA fragments (Rohland *et al*., 2018).

### Library Preparation

At the University of Tübingen, we used 25 µl of DNA to construct double-stranded, double-indexed libraries using a uracil-DNA-glycosylase treatment (UDGhalf protocol) (Rohland *et al*., 2015) Two indices per sample were selected by an in-house software to create a unique combination for each library. All libraries were initially PCR-amplified for 10 cycles and purified using MinElute columns with Qiagen reagents. Afterwards, libraries were re-amplified with library-specific cycles to aim for a concentration of 1.5E^13^ copies per library before purification. Afterwards, all libraries were quantified on the Agilent 4200 Tape Sstation System following the manufacturer’s protocol and pooled equimolar (10 nMol) for shallow shotgun sequencing. To increase nuclear coverage, we generated two additional single-stranded, double-indexed libraries for HELA002, following a non-Uracil-DNA glycolase protocol (Gansauge *et al*., 2020).

At the University of Pavia, 10-20 µl DNA was used to construct double stranded, double indexed libraries, implementing a half-Uracil-DNA glycolase (UDGhalf) protocol (Rohland *et al*., 2015). All libraries are PCR amplified for 12 cycles and purified using the QIAQuick MinElute purification kit (QIAGEN). The purified libraries are quantified using the Agilent 4150 TapeStation system and pooled equimolar for shallow shotgun sequencing.

At the Harvard medical school, the DNA was converted into sequenceable form using a single-stranded library preparation protocol, pretreated with uracil-DNA glycosylase (UDG) to minimize cytosine-to-thymine errors common in ancient DNA (Gansauge *et al*., 2020).

### Authentication, in-solution-capture and sequencing

To assess the presence of ancient DNA (aDNA) we considered the percentage of human DNA, the average fragment length and the damage pattern at the 5-prime end of the molecules. This screening process was performed with the software EAGER version 1.92.38 and integrated programs (Peltzer *et al*., 2016). The adapters were removed using AdapterRemoval with its default settings (Schubert *et al*., 2016). Afterwards, reads were aligned to the human reference genome hg19 (GRCH37) using the Burrows-Wheeler-Aligner (BWA) with disabled seedlength, a mismatch parameter of 0.01 and a quality filter of 30 (Li & Durbin, 2009). Afterwards duplicates were removed with DeDup and damage patterns were inferred using mapDamage2.0 (Jónsson *et al*., 2013). The selected samples were then re-amplified and enriched for a targeted set of 1.24 million single nucleotide polymorphisms (SNPs) across the human genome and the complete human mtDNA at the University of Tübigen and Harvard University (Table S1). Both target enrichments were performed in solution using a modified version of Fu et al (Fu *et al*., 2013). Single-end sequencing for samples captured at the University Tübingen was performed on an Illumina NovaSeq6000 instrument. Paired-end sequencing for samples captured at Harvard University was performed on an Illumina NextSeq500 instrument.

### Contamination estimates

Additional authentication steps were implemented on the captured data by estimating mtDNA as well as X-chromosome contamination levels. All results are detailed in Table S1. MtDNA contamination was established for single-stranded and double-stranded libraries separately using schmutzi and the appropriate settings for each library type (Renaud *et al*., 2015). X-chromosome contamination was estimated using Angsd (Korneliussen *et al*., 2014) for individuals genetically determined as males. For libraries with contamination levels over 5% (LAPE001, TPAG001, TPGU004), we performed PMD-filtering using PMDtools (Skoglund *et al*., 2014) with standard parameters (-threshold 3) for double-stranded libraries.

### Genotyping, Sexing and Haplogroup Calling

Before calling variants for double-stranded libraries, 2 basepairs (bp) at both reads termini were trimmed using the samtools trim package (https://github.com/samtools/samtools). For single-stranded libraries, no trimming occurred. Afterwards the pseudohaploid genotypes were called with a random calling approach implemented in pileupCaller, using the double-stranded and single-stranded options for each library (https://github.com/stschiff/sequenceTools) and later merged.

The biological sex was assessed using a custom script. The script estimates SNP coverage on the autosomes, X, and Y chromosomes, which is then used to calculate the Y and X ratios relative to autosomal coverage. MtDNA haplogroups were assigned on the consensus output files of schmutzi investigating quality scores between 10 and 30 using the software Haplogrep3 (https://haplogrep.i-med.ac.at/). Y-chromosome haplogroups were assigned from the bam files with Yleaf (https://github.com/genid/Yleaf) and Yhaplo (https://github.com/23andMe/yhaplo).

### Radiocarbon dating

We report new direct radiocarbon dates for 13 individuals (Table S1) performed at the Curt Engelhorn Centre for Archaeometry in Mannheim (Germany) and at the Centro di Datazione e Diagnostica of the University of Salento (Italy). At the dedicated facilities of the Curt Engelhorn Centre for Archaeometry, collagen was extracted using a modified Longin method and purified by ultrafiltration. Only fractions larger than 30 kD were retained. After purification, samples were freeze-dried and combusted to CO_2_ in an Element Analyser, catalytically converted to graphite, and AMS-analysed in a MICADAS-type machine simultaneously with a calibration standard (Ocalic Acid-II), blanks, and controls. All ^14^C-dates were normalised to δ^13^C+-25‰ and calibration occurred using the IntCal20 (Stuiver & Polach, 1977) dataset in combination with the SwissCal software (L.Wacker, ETH-Zürich). At the Centro di Datazione e Diagnostica (CEDAD) of the University of Salento, macro-contaminants were first identified by optical microscopy and mechanically removed, followed by chemical pretreatment using alternating acid-alkali-acid washes to eliminate remaining contaminants. The cleaned material was combusted at 900°C in an oxidizing environment to convert it into CO₂, which was subsequently reduced to graphite using H₂ and iron powder as a catalyst. Radiocarbon concentrations were measured by high-resolution accelerator mass spectrometry (AMS), with δ¹³C values determined directly by the accelerator to correct for isotopic fractionation. Standard reference materials, including sucrose C6 (IAEA) and oxalic acid (NIST), were used to ensure measurement accuracy. Conventional radiocarbon ages (BP) were then calibrated to calendar years using OxCal v3.10 and the INTCAL20 atmospheric dataset. Alternatively, we report previously published direct or indirect radiocarbon dates for other individuals in our dataset, when available (Table S1).

### Kinship, Runs of Homozygosity and Identity by Decent

We estimated biological kinship using the software KIN (Popli *et al*., 2023), specialised for low-coverage ancient data. We run KIN with standard parameters (-cnt 0, -r 1) and the KIN-estimated p-value. Due to a significant difference in baseline-relatedness between sites, questionable calls were repeated within sites. We estimated runs of homozygosity (ROHs), using hapROH for individuals with more than 400,000 SNPs covered on the 1240K panel (Ringbauer *et al*., 2021).

### Grouping and Labelling

To increase resolution in population-based analyses, we grouped individuals by site, considering available radiocarbon dates, archaeological information as well as genetic affinity (Tables S1 and S2). Due to clear differences in genetic affinity, AQUI005, LAPE001 and LAPE002 were treated as separate groups. The group labels were chosen to match the format: Site ID/Individual ID, average calibrated radiocarbon date in BP for each group.

### *f*-statistics and qpAdm analysis

For f-statistics we generated two different datasets: 1) individuals reported in this study intersected with the Allen Ancient DNA Resource (AADR) v62.0. (Mallick *et al*., 2024) 2) individuals reported in this study with present-day individuals genotyped on the Illumina panel (Reich *et al*., 2012). f_3_-outgroup statistics were computed using qp3Pop v.7.0.2 in the form f_3_(Mbuti; Pop1, Pop2). We also converted f_3_ values to dissimilarities by subtracting the values from 1 and creating MDS plots with a custom-made R script. A neighbour-joining tree was instead built with inverted f_3_-values (1/f_3_). *f*4 statistics were run using qpDstat v.7.0.2 with f4mode = YES (Patterson *et al*., 2012). We conducted qpAdm analyses using the Admixture tool (version 1.3.0) with standard parameters (maxrank=7).

#### Principal Component Analysis

For PCA construction we used the software smartpca (Patterson *et al*., 2006) with parameter *lsqproject:* YES. We used modern-day variation genotyped on the 1240K panel (Mallick *et al*., 2016) to build the PCA and projected ancient samples if they passed the threshold of at least 10,000 overlapping SNPs.

## Data Availability

The mtDNA and nuclear DNA sequences of the analyzed individuals in this study are available at the European Nucleotide Archive (ENA) under study accession number PRJEB102023 (data available upon publication). All data needed to evaluate the conclusions in the study are present in the main text and/or the Supplementary Information.

## Acknowledgements

We thank members of the Archaeo- and Palaeogenetics group at the University of Tübingen for comments and technical assistance. We thank Arturo Toscano and Carina Erchini (Museo Nacional de Antropología) for providing the samples from Uruguay and Mariano del Papa from Museo de La Plata who select and prepare the samples from Museo de La Plata. We are very grateful to Uruguayan odontologist Juan Pablo Oliver Espino for processing the Uruguayan samples and producing 3D-printed images to preserve the historical pieces. We thank INCUAPA-CONICET-UNICEN (Faculty of Social Sciences, National University of the Centre of the Province of Buenos Aires) for providing institutional support to carry out the sampling. We thank Claudia Della Negra (Direccion de Patrominio Cultural Material - Neuquen province) for the support provided during the sampling. This study received support from: the Italian Ministry for Universities and Research (MUR) for PRIN2022 2022Y8BSAL (to AA); Fondazione Cariplo - Bando Giovani Ricercatori 2023, rif: 2023–1373 (to NRM).

## Author contributions

K.L.K., performed the genetic analyses under the supervision of C.P. K.L.K., E.R., T.T., A.M.C.O., C.D., A.A., D.R., C.P. performed or supervised laboratory work. K.L.K., J.M.L., N.R.M., performed bioinformatics data processing. M.B.P., L.S., H.N., M.B., V.B., M.E.G., N.S., P.G.M., G.F., J.L.M., S.I.P., G.P., G.M., assembled archaeological material, described archaeological sites, and advised on archaeological background and interpretation. K.L.K., and C.P. wrote the manuscript, with inputs from all co-authors. J.L.M., S.I.P., G.P., G.M., C.P. initiated and supervised the study.

