## Supplementary Information for "The shared genomic history of Middle to Late Holocene Southern Cone populations"

Archaeological site descriptions

We report newly generated genomes from 52 ancient individuals, from 29 sites and three regions of Argentina, namely the central and Southern Pampas, Northwest Patagonia and the Paraná River Delta, as well as two sites from two regions in Uruguay, i.e., the Lower Uruguay River and the eastern lowlands of Uruguay.

### Central Pampas

#### La Macarena (LAMA_3900BP)

I20341 (1B): 3668 ± 18 BP

#### Las Tunas Chicas (LATC_3100BP)

I20342 (2b): 2965 ± 15 BP

#### Zig-Zag (ZIZA_1,900BP)

I20197 (6A): 2020 ± 20 BP

#### Beruti (BERU_1500BP)

I20346 (7B): 1608 ± 15 BP

#### Loma Alta (LOAL_600BP)

I20345 (5B): 630 ± 11 BP

These five inhumations are located in the Hinojo-Las Tunas Shallow Lake System, one of the largest permanent lake cluster in the Central Pampean Dunefields area of the Central Pampas (La Macarena: -35.984333/-62.333806; Las Tunas Chicas: -36.075792/-62.363908; Loma Alta: -35.903753/-62.47315; Beruti: -35.849753/-62.461797; Zig-Zag: -35.877136/-62.473375). Forensic police recovered all the individuals after they were exposed to the surface due to the advance and retreat of the shallow lake margins, which caused erosion of the adjacent dunes. All correspond to simple primary burials represented by adult individuals. Chronologically, the individuals were radiocarbon dated at various points in the Late Holocene (Scheifler *et al.*, 2024)(Table S1). The remains from these burials are deposited in the bioarchaeological collection of the Museo Histórico Regional de Trenque Lauquen.

### Southern Pampas

#### Tres Bonetes 1 (TREB_5200 BP)
#### Cantera de Rodados Villalonga (CANT_4800BP)

TREB001 (FCS.TB.E1.1): 5188 ± 40 BP
TREB002 (FCS.TB.E3.1): 5339 ± 39 BP
CANT001 (FCS.CRV.E2.1): 4889 ± 58 BP

These sites are located in the southern Pampas (Pampa-Patagonia transition) close to each other, along the Atlantic coast (-40.100028°/62.350083° (CANT); -40.190361°/-62.363000° (TREB)). In both contexts, human remains were recovered on marine deposits in stratigraphic position. From the seven individuals recovered, six belong to amateur collections and one was excavated. Chronology ranges from 5300 to 4100 years BP. Inhumations correspond to the primary burial modality and these contexts have been interpreted as human burial settings. No grave goods were recorded in association with the burials (Martínez *et al.*, 2024).

**Paso Alsina 1 (ALSI_480BP)**

ALSI001 (FCS.PA1.E2.I36)
ALSI004 (FCS.PA1.E4.I46)
ALSI006 (FCS.PA1.E3.I41)
ALSI007 (FCS.PA1.E8.I48)
ALSI008 (FCS.PA1.E2.I29)
ALSI009 (FCS.PA1.E6.I55)
ALSI010 (FCS.PA1.E3.I39)
ALSI012 (FCS.PA1.E10.I8)
ALSI013 (FCS.PA1.E1.I31)

The Paso Alsina 1 site (Southern Pampas/Pampa-Patagonia transition) is located 400 m from the right bank of the Colorado River and 100 km from the Atlantic coast (-39.390833°/-63.260000°). A total of 10 multiple secondary burials were recovered from a small area (6m$^2$). The secondary burials follow a spatial arrangement produced by an intentional mortuary practice of patterned orientation and contiguous and/or overlapping arrangement of bundles. The burials are distributed only over a vertical range of ca. 30-40 cm.

Six bundles were aligned in an east-west direction, and the remaining four in a north-south orientation. The mortuary context does not present any evidence of modification resulting from the inclusion of further interments, indicating that the bundles were made, transported, and deposited at this particular place in the landscape without further human disturbance or the removal of bones. Thirteen radiocarbon dates from all bundles yielded values consistent with one another, yielding a weighted average of 483 ± 20 years BP. Based on this chronological pattern and the site's contextual features, we can infer that individuals died over a brief period and that bundles were buried synchronously (Martínez *et al.*, 2012).

A minimal number of 77 individuals (MNI) was estimated (Flensborg *et al.*, 2015). The spatial arrangement of the anatomical units within the bundles showed a similarity in the association of bone elements, allowing a so-called “basic structure” to be defined. The basic structure consists of one to three skulls on the ends, bones of the pelvic and shoulder girdle associated with the skulls, long bones on the sides, and groups of ribs aligned symmetrically with each other on top of the bundles (Martínez *et al.*, 2012). Despite this formal pattern, each bundle presented differences with respect to the location and quantity of bone elements (e.g., more than three skulls on the ends, skulls in the middle of the bundle, etc. (Martínez *et al.*, 2006).

An interesting aspect to note is that most of the bones show no anatomical articulation with each other, except for some skulls and jaws and a few dorsal vertebrae. In this regard, the osteological record suggests that the bundles were made with individual anatomical units (Martínez *et al.*, 2006). The clear evidence of manipulation of corpses is further reinforced by cases like that of a 2 to 3 year-old infant's skull intentionally disarticulated at the sutures, the bones of which (parietal, frontal and occipital) were placed on top of each other and, in turn, assembled on top of an adult female’s pelvis (Martínez *et al.*, 2012).

The sex-age analyses indicate that individuals of both sexes and different age categories (fetuses, infants, children, adolescents and adults) are present in the funerary structure (Flensborg *et al.*, 2015). Nonetheless, every bundle presents a differing age-sex structure:

some burials (e.g., burials 4, 8 and 9) are composed of a larger number of sub-adult individuals, while others (e.g., burials 2, 3 and 5) are mostly represented by adults. Regarding traces of body manipulation, evidence of defleshing, cutting and scraping have been recorded (Martínez *et al.*, 2012). Despite the presence of these anthropogenic marks a significant number of skeletal elements do not show such modifications, suggesting that bones were in various stages of natural decomposition and skeletonization. A remarkable aspect of the mortuary practice is the intensity of the application of colour to all bones, regardless of their position within the bundle. Red dye or paint is present on all bone surfaces. Furthermore, the uniformity of the colour and the staining of inner portions of some skulls and long bones suggest that the substance was liquid enough to reach even the internal portions of anatomical units.

As at the other sites in the area with human burials, there were no personal ornaments or funerary goods associated with individuals in the bundles or funerary structure. The site was therefore defined as an exclusive area of inhumation that consists solely of multiple secondary burials deposited in a single event.

**El Remo (REMO_230 BP)**

REMO001 (FCS.ER.E1.1): 230 ± 42 BP

The El Remo site  (Southern Pampas/Pampa-Patagonia transition) is located 2 km from the right bank of the Colorado river and approximately 20 km from its mouth (-39.747528°/-62.369722°). A single primary burial, female and adult, that comes from an amateur collection was dated at ca. 250 years BP. Along with the skeletal remains eight glass beads were identified. No functionality of this site was possible to assigned due to the scarcity of data provided from the collector (Flensborg & Wagner, 2015).

**La Primavera (PRIM_2700BP)**

PRIM001 (FCS.PRI.E1.1) and PRIM002 (FCS.PRI.E1.1d): 2728 ± 48 BP

La Primavera site (Southern Pampas/Pampa-Patagonia transition) is located 5 km from the left bank of the Colorado river and approximately 10 km from its mouth (-39.590833°/ -62.391389°). Two incomplete primary burials were recovered and dated at ca. 2900-2700 years BP. The individuals were adult males. One of the skeletons was on its side, legs flexed. No grave goods were recorded in association with the burial except for some possible personal ornaments, including an *Amiantis purpurata* shell. The archaeological material recovered at the site (lithic artifacts, faunal remains, etc.) led to the site's characterization as a multi-activity residential base where burials also took place (Bayón *et al.*, 2004).

**La Petrona (LAPE001_300BP, LAPE002_400BP)**

LAPE001 (FCS.LP.E1.1): 352 ± 51 BP
LAPE002 (FCS.LP.E3.1): 436 ± 39 BP

This site (Southern Pampas/Pampa-Patagonia transition) is located on a prominent sand dune, 200 m from the north bank of the Colorado River and ca. 50 km from the Atlantic coast (-39.503867°/ -62.785553°). Two primary and two multiple secondary burials were recovered in an area of 25 $m^2$ (Martínez & Figuerero Torres, 2000).

One multiple secondary burial (LP1) is composed of three adults, two women and one individual of undetermined sex. The presence and frequency of skeletal parts indicates that individuals are represented by incomplete skeletons. There is a pattern in the bone distribution: the bundle consists of a skull and pelvises at each end, long bones at the sides, and other bones (vertebrae, ribs, bones of the hands and feet) in the central part. Bones

from the burial lack any kind of anatomical articulation. Nevertheless, regularities in the disposition of some bone elements belonging to the same individual within the bundle were recorded. A close spatial association of bones anatomically related (e.g., humerus and radius), as well as associations between homologous bones (e.g., left and right femur), was recognized (Martínez & Figuerero Torres, 2000). Finally, bone elements of the axial skeleton, hands, and feet were found inside the central part of the bundle. All these features correspond to the intentional manipulation of the anatomical units in the bundle assemblage. Two radiocarbon dates indicate a chronology of 352 ± 51 years BP and 314 ± 45 years BP (Flensborg *et al.*, 2011).
The other secondary burial (LP2) is also composed of multiple, incomplete skeletal parts of three individuals, one adult female of unknown age, and an infant. The organization of bones within this bundle resembles LP1. There were no anatomically articulated bones. However, some bones belonging to the same individual were found close together (e.g., scapula and humerus). Sediments with red pigment were present in the center of the bundle (Martínez & Figuerero Torres, 2000). A small number of bone elements with evidence of red colouring on the surface was recorded. Regarding the manipulation of corpses, there were small and faint traces of scraping and cutting related to body disarticulation and bone surface cleaning (Flensborg *et al.*, 2011). Two radiocarbon dates obtained from two bones from this bundle place the burial between 481 ± 37 and 770 ± 49 years BP). Due to the fact that this bundle is a multiple secondary burial and the chronological data obtained, it is proposed that the two dates correspond to different individuals, suggesting that human groups were gathering up the bones of individuals who died at different times and, probably, at different places in the landscape in order to perform secondary burials (Flensborg *et al.*, 2011).
A further primary burial (LP3) is composed of an incomplete skeleton of an adult female located on her side. The particularity of this burial is the absence of some of the anatomical units of a fully articulated skeleton: the bones from the third lumbar vertebrae down are absent, as well as the rest of the bones of the spine, the pelvic girdle, and the lower members. Interestingly, the clavicles and two thoracic vertebrae that occupy intermediate positions between other thoracic elements are also missing (Flensborg *et al.*, 2011). Analysis indicates that this pattern cannot be attributed to the action of taphonomic or post-depositional processes. Instead, the absence of the anatomical units in this burial has been explained by the intentional removal and recovery of bones for the making of bundles (Martínez & Figuerero Torres, 2000; Flensborg *et al.*, 2011). Two radiocarbon dates indicate a chronology of 411 ± 39 and 462 ± 39 years BP.
The remaining primary burial (LP4) is composed of an incomplete skeleton pertaining to an adult female individual. Unlike the other burials, the anatomical units were mainly found scattered -- only a few elements remained articulated. Taphonomic processes appear to have played an important role in the pattern of bone preservation and distribution (Flensborg *et al.*, 2011). The remaining articulated elements were part of the pelvis and bones of the lower limbs, which were flexed. On some of the elements (e.g., pelvis and tibia) red mottling suggests the presence of mineral pigments (Martínez & Figuerero Torres, 2000). There were traces of body manipulation such as defleshing and scraping. Unlike the secondary burial LP2, marks were more frequent and visible, sometimes grouped in clusters, and indicate that soft tissue was removed before primary burial. Therefore, the body did not undergo a process of natural skeletonization as a result, for example, of primary burial. Rather, the skeleton was intentionally defleshed to extract the soft elements, which accelerated the decomposition of the body and possibly the disarticulation of the bones to facilitate their transport to other parts of the landscape (Flensborg *et al.*, 2011). A radiocarbon date of 248 ± 39 years BP was obtained.
No grave goods or personal adornments were found accompanying these burials. The dune where the burials were deposited has a surface distribution of many different types of artifacts. Among those, a variety of types of projectile points and lithic debris, grinding tools, lip and ear ornaments, faunal remains, and pottery sherds were recorded. The characteristics of the archaeological record of La Petrona site enable it to be categorized as

a multi-activity residential base where funerary practices were also conducted, including the handling of corpses and skeletons (Martínez & Figuerero Torres, 2000).

### Northwest Patagonia

#### Hermanos Lazcano (HELA_4300BP)

HELA002 (HL II-est.5-cud 1A): 3780 ± 50 BP

Hermanos Lazcano is an open-air site located in an alluvial sediments deposit in the Chacay Melehue valley (-37.373° / -70.368°), 5 km from El Alamito in Neuquén province. The site dates back between 4100 and 4450 years Cal BP (Cobos *et al.*, 2022). The archaeological site comprises multiple burials of at least 15 individuals. These are adult and subadult individuals of both sexes arranged in primary inhumations (Della Negra *et al.*, 2014). The archaeological material associated with the burial consists of stones grinding tools, personal adornments made of valves, lithic instruments, and rectangular flat stones that mark the burial sites and are generally located near skull (Della Negra & Saint Paul, 2012; Della Negra *et al.*, 2014).

#### Aquihueco (AQUI_4200BP, AQUI005_4200BP)

AQUI003 (Aquihueco 1 ind 33): 4045 ± 66 BP
AQUI005 (Aquihueco conj 2 ind 1)
AQUI007 (Aquihueco 1 ind 38)

Aquihueco is an open-air archaeological site located on a sand dune in the Curi Leuvu valley (-37.093 / -70.377), 8 km from Tricao Malal, in Neuquén province. The site dates back between 3930 and 4835 Cal year BP (Cobos *et al.*, 2022). The archaeological site corresponds to multiple burials of a minimum number of 64 individuals. There are adult and subadult individuals of both sexes arranged in primary inhumations (Della Negra & Novellino, 2005; Della Negra *et al.*, 2006; Gordón *et al.*, 2019). The structural characteristics of the burials include the demarcation of the inhumations by clusters of small stones, large flat or columnar rocks, while other burials had no demarcation. In direct association with some adults and infants, personal adornments –malacological and rock pieces– of symbolic value were found arranged near the cervical vertebrae, sternum, and skull (Della Negra & Saint Paul, 2012). Analysis of bone and dental tissues of the individuals indicated high degrees of dental wear and a low frequency of caries (Della Negra & Novellino, 2005; Gordón *et al.*, 2019). Several individuals had a type of cultural modification of the skull known as pseudo-circular (Perez *et al.*, 2009).

#### Sitio Grande (GRAN_600BP)

GRAN001 (Grande): 670 ± 40 BP

Sitio Grande is an open-air archaeological site located on an island in the Limay river (-39.65° / -69.30°), 19 km from Picun Leufu locality in Neuquén province. The site dates back 605 years cal BP (Cobos *et al.*, 2022). The archaeological site corresponds to multiple burials of at least 3 individuals. These are adult and subadult males arranged in primary burials (Novellino, 2002). The associated archaeological material consists of ceramic pottery fragments and lithic materials (Novellino, 2002).

#### Loma de la Lata (LOLA_600BP)

LOLA001 (Loma de la lata 7)

Loma de La Lata is an open-air site located in Cerro de los Indiecitos, close to Neuquén river (-38.417° / -68.683°), 14 km from Añelo locality in Neuquén province. The burial site dates back between 640 and 560 years Cal BP (Cobos *et al.*, 2022). The archaeological site corresponds to multiple burials of at least 19 individuals. These are adults and subadults individuals of both sexes arranged in primary and secondary burials (Cúneo *et al.*, 2016). The burials were found covered with flagstone. The archaeological material associated are triangular projectile points, tembetás, perforated shells, necklaces made from snail shells, beads made from mollusk shells, and blue-green rocks containing copper minerals (possibly chrysocolla; (Cúneo *et al.*, 2016). Most individuals present planolambdic cranial modification (Perez *et al.*, 2009).

**Calle Lanin (CALA_500BP)**

CALA001 (Calle Lanin 1): 385 ± 16 BP

Calle Lanin is an open-air site located near Limay valley (-38.9647° / -68.0727°), in Neuquén city of Neuquén province. The site dates back to historical moments in 390 years Cal BP. The archaeological site corresponds to a simple burial of primary inhumation of an adult female deposited with lower limbs flexed. No funerary structure, grave goods or personal adornments were found accompanying the burial. No present intentional artificial modification of the skull.

**Campo Ayoso (CAYO_500BP)**

CAYO001 (Campo Ayoso 1): 385 ± 16 BP

Campo Ayoso is an open-air archaeological site located on Alumine river valley (-39.2329° / -70.9234°), in Alumine city in Neuquén province. The site is dated in the 390 years Cal BP. It is a primary multiple burial of two adult females. The burial did not contain any associated archaeological material.

**Chacra Bustamante (CABU_460BP)**

CABU001 (Chacra Busamante): 450 ± 40 BP

Chacra Bustamante is an open-air archaeological site located in a field (-39.05° / -69.05°), 18 km from Cutral-co locality in Neuquén province. The site is dated in 470 years Cal BP (Cobos *et al.*, 2022). It corresponds to the burial of an adult male arranged in primary inhumations (Perez *et al.*, 2009). The archaeological material associated are small projectile points with triangular limbs and notched and straight bases, bone instruments, decorated ceramics, scrapers and grinding instruments (Perez *et al.*, 2009).

**Remeco (REME_290BP)**

REME002 (Remeco I ind. 4): 230 ± 40 BP

Remeco is an open-air archaeological site located in Remeco stream valley (-39.0742° / -71.3276°), 17 km from Moquehue locality in Neuquén province. The site is dated in 195 years Cal BP (Moscardi *et al.*, 2024). It corresponds to multiple burials of a minimum number of 5 individuals (Bernal *et al.*, 2021). These are adult and subadult individuals arranged in primary inhumations in three funerary structures. Each of these had a lid, walls and base built with flagstone, they have been called cists by Hajduk (1981-82), and were oriented in a N-S or N-NE/S-SW direction. The archaeological material associated with the burial consists of fauna bone remains, ceramic and lithic materials, metal elements, mollusc shells, textiles, glass necklace beads and leather (Bernal *et al.*, 2021).

**Cape Malal (CAMA_200BP)**

CAMA001 (Caepe malal E4): 200 ± 50 BP

Cape Malal is an open-air site located on Curí Leuvú River valley (-37.1655° / -70.3977°), 32 km from Chos Malal city in Neuquén province. The site is dated in the 18th-century (Hajduk & Biset, 1991; Hajduk *et al.*, 1996). The archaeological site corresponds to multiple burials of a minimum number of 12 individuals. They are adult and subadult individuals of both sexes arranged in primary inhumations. Among the archaeological materials found were copper and iron objects knives, bells, buckles, needles and buttons–, glass beads, pottery, fragments of leather from a helmet, an iron saber, and a boleadora ball. In addition, faunal remains of horse, dog, mollusks from the Pacific and ovicaprids were found (Hajduk *et al.*, 2000). Most individuals present planolambdic cranial modification (Perez *et al.*, 2009).

**Piera (PIRA_195BP)**

PIRA001 (Piera): 229 ± 32 BP

Piera is an open-air site located on Covunco stream valley (-38.703° / -69.983°), 7 km from Mariano Moreno city in Neuquén province. The site dates back to 195 years Cal BP (Gordón *et al.*, 2019; Cobos *et al.*, 2022). This burial consists of a genetically female adult, buried in an extended position. Among the archaeological materials found were silica and obsidian artifacts, a fragmented boleadora ball, as well as metal fragments and a glass bead. A set of sedimentary rocks were placed on top of the burial, possibly for covering the individual.

**Andacallo (ANDA_160BP)**

ANDA001 (Andacallo): 168 ± 18 BP

Andacollo is an open-air site located on Neuquén river valley (-37.1764° / -70.6661°), in Andacollo city in Neuquén province. The site dates back 109 years Cal BP. It corresponds to primary inhumations of two individuals, an adult male and a subadult, associated with ancient European material The adult individual presents a planolambdic cranial modification. Among the archaeological materials found were pottery and iron elements for horses. In addition, faunal remains of a horse were found.

**Añelo (ANLO_historical)**

ANLO001 (Añelo 0001): 200 ± 50 BP

This is an individual probably coming from the Loma de La Lata site (LOLA) dated by association to the historical period.

**Alonqueo (ALON_historical)**

ALON001 (Alonqueo I 3): 200 ± 50 BP

Alonqueo 1 is an open-air site located on Agrio river valley (-38.5490° / -70.3726°), 3 km from Mariano Moreno city in Neuquén province. The site is dated in the 172 years Cal BP. It is a simple primary burial, with old European material associated, as well as faunal remains of horses.

### Paraná River Delta

#### Laguna de los Gansos 2 (LAGA_570BP)

LAGA001 (LDLG2-1,Arg006): 570 ± 43 BP
LAGA002 (LDLG2-2, Arg007): 590 ± 46 BP

The site Laguna de los Gansos 2 (32º29′37.1″ S, 60º38′25.7″ W) was studied by Bonomo and Politis between 2012 and 2014 (Bonomo *et al.*, 2016). It is located on a natural levee at the entrance of a corral adjacent to a rural outpost in the Diamante Department, Province of Entre Ríos. A total area of 17 m² was excavated. The excavations yielded 587 ceramic fragments—including plain sherds, incised pieces, vessels decorated with red pigment, and zoomorphic appendages—along with 461 faunal remains. Among mammals, the most represented taxa were *Myocastor coypus*, *Hydrochoerus hydrochaeris*, and *Blastocerus dichotomus*. Additional remains included *Actinopterygii* and *Siluriformes* fishes, as well as *Mollusca*. Human skeletal remains from two individuals were also recovered: one from a primary burial and another from a secondary burial, both dated between 570 and 590 $^{14}$C years BP (Bonomo *et al.*, 2016).

The specimen analysed in this study (LAGA001) corresponds to the right petrous bone of an adult male. This incomplete skeleton, found in a dorsal decubitus position, preserved the pelvic girdle with articulated lower limbs, and the skull was relocated over the pelvis, while the remaining bones were absent. The individual was dated to 570 ± 43 $^{14}$C years BP (AA-98851), with the following isotopic values: $\delta^{13}$C −19.9‰, $\delta^{13}$ = −14.4‰, and $\delta^{15}$N = 10.2‰ (UGAMS-11476). The second specimen (LAGA002) corresponds to the left petrous bone of an adult individual, dated to 590 ± 46 $^{14}$C years BP (AA-103899), with an isotopic value of $\delta^{13}$C = −20.2‰ (Bonomo *et al.*, 2017).

#### Cerro de las Pajas Blancas 1 (CPAB_175-750BP)

CPAB001 (CDLPB1-E1, Arg009): 175 ± 45 BP
CPAB002 (CDLPB1-E2-35, Arg010): 754 ± 40 BP
CPAB003 (CDLPB1-E2-4, Arg011): 720 ± 40 BP

Cerro de las Pajas Blancas 1 (CDLPB1) (32°6′36.8″ S, 60°44′33″ W) is located on La Vencida Island in the lower Paraná River (San Jerónimo Department, Santa Fe Province, Argentina). The site lies on a 179 m long, 4 m high levee adjacent to Cerro Madrejón. CDLPB1 has long been a key reference in regional archaeology, known since the 1940s for the discovery of Guaraní pottery, including a polychrome urn containing human remains. Based on this find, Serrano (1950, 1955) (Serrano, 1950; 1955) defined the “Pajas Blancas Polychrome” ceramic type.

Early studies (Badano, 1940; Serrano, 1955; Gollán, 1989) debated the association between Guaraní materials and local Chaná-Timbú traditions, interpreting the site as an isolated Guaraní settlement (Caggiano, 1983) linked to their southern expansion. After more than three decades without new excavations, research resumed in 2006–2007 (Bonomo *et al.*, 2011b), including mapping, surface collections, a stratigraphic test pit, and radiocarbon dates (ca. 640–650 $^{14}$C years BP). Analyses recorded plain, incised, painted, and corrugated pottery. The study of the faunal remains identified coypu, fish, and freshwater mollusks. Starch grains of *Phaseolus* and *Zea mays* were also identified (Bonomo *et al.*, 2011a). Further research (Sartori, 2013; 2015) added ceramic and zooarchaeological data, as well as another date of ca. 500 $^{14}$C years BP.

New systematic excavations in 2018–2019 expanded the dataset with 15 m² opened, yielding over 2,500 ceramic sherds, a nearly complete globular vessel, personal ornaments (earspools, a T-shaped tembetá, a jaguar canine pendant, etc.), faunal and malacological remains, and human burials. Microbotanical analyses identified the presence of both wild and domesticated plants, including maize phytoliths.

The CDLPB1 assemblage is regionally significant because it integrates elements of the two major archaeological entities in the study area: Goya-Malabrigo (Serrano, 1950; Politis & Bonomo, 2018; Politis & Bonomo, 2023) and Guaraní (Torino *et al.*, 2023). Goya-Malabrigo components are represented by rhythmic-incised pottery, bi- and tridimensional zoomorphic appendages, ceramic bells, intensive exploitation of aquatic fauna, and possible anthropogenic elevation of the levee. Guaraní materials occur in smaller proportions, including corrugated, fingernail-impressed, and polychrome ceramics, as well as a diagnostic urn burial (Torino *et al.*, 2023).

At least six human individuals—both adults and subadults of different ages and sexes—were interred at CDLPB1, indicating that the site was not restricted to a particular demographic group. Samples analyzed include the right petrous bone of a young adult male (CPAB001) from a primary burial accompanied by a rich funerary assemblage: two bone tools (on a fish rib and a cervid metapodial), a ceramic earplug, a pendant made from the right upper canine of *Panthera onca* (perforated at the root, longitudinally split, and polished), a left lower canine of *Pecari tajacu*, and a *Diplodon* sp. shell. Additional samples comprise the right petrous bones of a subadult individual (CPAB002) and an adult individual (CPAB003). These remains had been exhumed and subsequently reburied by local rural inhabitants, resulting in the complete loss of their original contextual association.

Besides the outlier essay of 175 ± 45 $^{14}$C years BP (LTL21136C) obtained for the primary burial (CPAB001), which we rejected, two other individuals yielded statistically indistinguishable dates of 754 ± 40 $^{14}$C years BP(LTL21137C, CPAB002) and 720 ± 30 $^{14}$C years BP (LTL21138C, CPAB003).

**Túmulo I del Paraná Guazú (TPGU_770BP)**

TPGU001 (MLP-DA-30, Arg030): 777 ± 40 BP
TPGU002 (MLP-DA-32, Arg032)
TPGU003 (MLP-DA-39, Arg039)
TPGU004 (MLP-DA-46, Arg046)

Túmulo I del Paraná Guazú (S34°1′, W58°41′) was first investigated by Torres in 1905 (Torres, 1911) and later by Lothrop in 1925 (Lothrop, 1931), who renamed it *El Cerrillo*. The mound is located on a levee beside a small stream flowing into the right bank of the Paraná Guazú River, between the Paraná Miní course and the Segunda Campana channel (Campana District, Buenos Aires Province). Based on the reports of Torres (Torres, 1911) and Lothrop (Lothrop, 1931), the combined excavations covered an estimated area of 755 m² and yielded more than 60 human skeletons. Materials from the site are curated at the La Plata Museum, Argentina (MLP), and the National Museum of the American Indian, USA (NMAI).

Reanalysis of these collections (Bonomo *et al.*, 2009; Bonomo, 2013; Ramos *et al.*, 2018) revealed a predominance of ceramics, including 1,074 sherds (plain, incised, and corrugated), a small bowl (7.7 cm in height and 7.5 cm in rim diameter), three clay lumps, and an unfired clay roll. Other finds include four copper sheets, 95 lithic artifacts (bola stones, mortars, grinding stones, hammers, anvils, a projectile point, and unmodified pebbles), and 260 bone objects, of which 111 are bone tools (conical points, harpoons, bevels, tubes, and perforated cervid forks). Faunal remains identified at the site comprise bird bones (undetermined taxa), cervids (*Blastocerus dichotomus*, *Ozotoceros bezoarticus*, *Mazama* sp.), siluriform fishes, ampullariid shells (*Pomella megastoma*), freshwater clams (*Diplodon parallelopipedon* and *Diplodon aff. variabilis*), and charred endocarps of the pindó palm (*Syagrus romanzoffiana*).

Regarding human remains, the reanalyzed assemblage (Ramos van Raap & Bonomo, 2016) comprises 241 specimens, from which a MNI=19 was estimated for Túmulo I del Paraná

Guazú and 9 for El Cerrillo. An AMS radiocarbon date obtained from a human bone at El Cerrillo yielded an age of 576 ± 42 $^{14}$C years BP (AA-93215) (Bonomo *et al.*, 2011a). The specimens examined in this study consist of temporal bone fragments with adhered petrous portions: one left (TPGU001) and three right (TPGU002, TPGU003, and TPGU004). TPGU001 was directly dated to 777 ± 40 $^{14}$C years BP (LTL21139C).

**Túmulo II del Paraná Guazú (TPAG_670BP)**

TPAG001 (MLP-DA-51, Arg051)
TPAG002 (MLP-DA-53, Arg053)
TPAG003 (MLP-DA-61, Arg061): 678 ± 35 BP
TPAG004 (MLP-DA-63, Arg063)

Túmulo II of Paraná Guazú (S 33°57′, W 58°47′), excavated and studied by Luis María Torres in 1911, is a circular mound located about 1,500 m from the confluence of the Carabelitas Stream with the Paraná Guazú River. The collection, curated at the Museo de La Plata and later reexamined (Bonomo *et al.*, 2009), comprises 45 ceramic sherds, 12 lithic tools, 6 lithic flakes, 5 bone tools, and 4 copper fragments. In addition to these domestic remains, the site also included a designated cemetery area, from which a minimum of 60 individuals were recovered (see Ramos *et al.*, 2018). A previous radiocarbon date obtained from a human bone yielded 846 ± 41 $^{14}$C years BP (AA-72633; (Bernal, 2008)). The samples analyzed in this study consist of temporal bone fragments with adhered petrous portions: three right (TPAG001, TPAG002, TPAG003) and one left (TPAG004). Sample TPAG003 was dated at 678 ± 35 $^{14}$C years BP (LTL21140C).

**Túmulo del Brazo Largo (TBLG_650BP)**

TBLG001 (MLP-DA-120, Arg120): 656 ± 42 BP

**T**úmulo I of Brazo Largo, excavated and studied by Luis María Torres (Torres, 1911), is a mound located on a levee adjacent to a small stream flowing into the left bank of the lower course of the Brazo Largo Creek (Islas del Ibicuy Department, Entre Ríos Province). The associated collection, curated at the Museo de La Plata and later re-examined (Bonomo *et al.*, 2009) includes 70 lithic artifacts (tools, cores, and debitage), 11 bone tools, 33 plain and decorated pottery sherds, and faunal remains such as freshwater mollusks (Ampullariidae, *Diplodon* aff. *variabilis*), indeterminate birds, and marsh deer (*Blastocerus dichotomus*). Starch grain analysis conducted on a grinding stone identified remains of *Phaseolus* sp., *Prosopis* cf. *nigra*, *Manihot esculenta?*, and *Zea mays* (Bonomo *et al.*, 2011b). At least seven individuals were recovered from this site, including a left temporal bone (TBLG001) that was analyzed in this study. From this same human sample, a radiocarbon date of 656 ± 42 $^{14}$C years BP (AA-93217) was previously obtained (Bonomo *et al.*, 2011a).

**<u>Eastern lowlands of Uruguay</u>**

**Los Indios (INDI_870 BP)**

INDI001 (LOS_INDIOS_EXCII_num12): 873 ± 20 BP

Los Indios archaeological site (UTM 33°53´57´´/53°42´42´´ W) is located in a strategic place for crossing extensive lowlands to the north of Negra Lagoon in the east of Uruguay. The site occupies both shores of the Los Indios stream. The research (1998 to 2011) focused on a multicomponent east shore of the site, which was occupied since at least 8500 to 7000 B.P as seasonal hunter camps to capture faunal from lowlands (deer, rodents and fishes). These old archaeological levels have accumulations of lithic material as grinding stones, many stemmed projectile points, scrapers, and other bifacial tools mostly made in siliceous raw material from at least 400 km of the north (López Mazz & Gianotti, 2001). Between 7,000 and 3,100 BP there is a gap in human occupation.

Later, between 3,000 BP and 700 BP, the site was transformed to a semi-circular village open to the east, with 3 mounds, earthwork, activity areas and one platform (López Mazz & Gianotti, 2001). This period shows the process of social intensification with mortuary practices in different modalities, suggesting social differences between (Gianotti & López Mazz, 2009). In this period, the economy continued the exploitation of deer (*Ozotoceros bezoarticus, Odocoileus dichotomus*), rodents (*Myocastor coypus, Cavia sp.*), and fish (*Siluriformes, sp.)* from the lowlands and sea mammals from the Atlantic coast (*Otaria flavescens*) as far as 15 km away (Moreno Rudolph, 2014).

Lithic technology changes in this period to more regional provisioning of raw material. Projectile points, in various shapes and propulsion systems, continue to be used with stone balls (boleadoras) and play a central role in hunter strategies (Clemente-Conte & Mazz, 2023). Very simple ceramics without decoration are also characteristic of these Cerritos Builder archaeological traditions of the east of Uruguay and south of Brazil (López Mazz, 2024). The economy of these groups was complemented with wild palms (*Butia odorata*) and domestic plants (*Zea mays* and *Phaseolus spp.*) (Iriarte *et al.*, 2001).

The excavations recovered human remains of 13 individuals (NMI) from mounds I, II, III and IV. One of these is a secondary bundle burial, five others are flex primary burials, and seven are clusters of bone fragments (Gianotti & López Mazz, 2009). Associated with the burials were recovered stone balls, lithic flakes and cores, and fragments of white quartz. The study of isolated human bones shows cut marks and thermal alteration interpreted as episodes of violence, scalping and cannibalism (Gianotti & López Mazz, 2009) in the context of significant social changes (López Mazz *et al.*, 2014). Mound II was built around 2800 BP and shows archaeological floors on fragmented human bones of one adult individual, accompanied by a stone polish ball. Sample INDI 001 is a tooth from an adult dated by radiocarbon in C14 Age (yr BP) 873+-20 (Calibrated Age (95%) AD.1053-1223).

### <u>Lower Uruguay River</u>

#### Los Cardos (LOCA_1700BP)

LOCA001 (LC1): 1,680 ± 17

Los Cardos site (UTM 33°28´34¨ S / 58°24´33´´W) is a multicomponent open-air site located on the shore of the Uruguay River, near the mouth of the San Salvador River. Research began in 1986 and has identified archaeological layers containing burials associated with abundant faunal remains, indicating economic practices focused on fluvial resources (Toscano, 1992).

The cultural material is similar to that of the nearby Cañada Saldaña and San Salvador sites, both of which belong to the Goya Malabrigo archaeological tradition. This tradition is characterised by primary extended burials, highly decorated ceramics, and complex bone technology used for fishing and hunting (López Mazz, 2018; Politis & Bonomo, 2018).

This region witnessed the beginning of the Spanish conquest of the Río de la Plata with the expeditions of Sebastián Gaboto (1527) and Ortíz de Zárate (1574), which had a significant impact on the endurance of Indigenous peoples (López Mazz *et al.*, 2014). Archaeological and ethnohistorical studies suggest cultural continuity between the Goya Malabrigo archaeological entity and the Chaná-Timbú tribes (Acosta y Lara, 1956; Politis & Bonomo, 2018). Archaeological excavations recovered three burials: two bundles containing bones from multiple individuals (n=3 and n=4 , respectively) and one primary extended burial (Erchini, 2000). Burial DOL1E2B, which was fragmented and poorly preserved, was associated with a ceramic pendant typical of the Goya Malabrigo tradition. Sample LOCA001 from this burial is a petrous bone from an adult, dated by radiocarbon (C14) to cal AD 263-418.

### Supplementary Figures

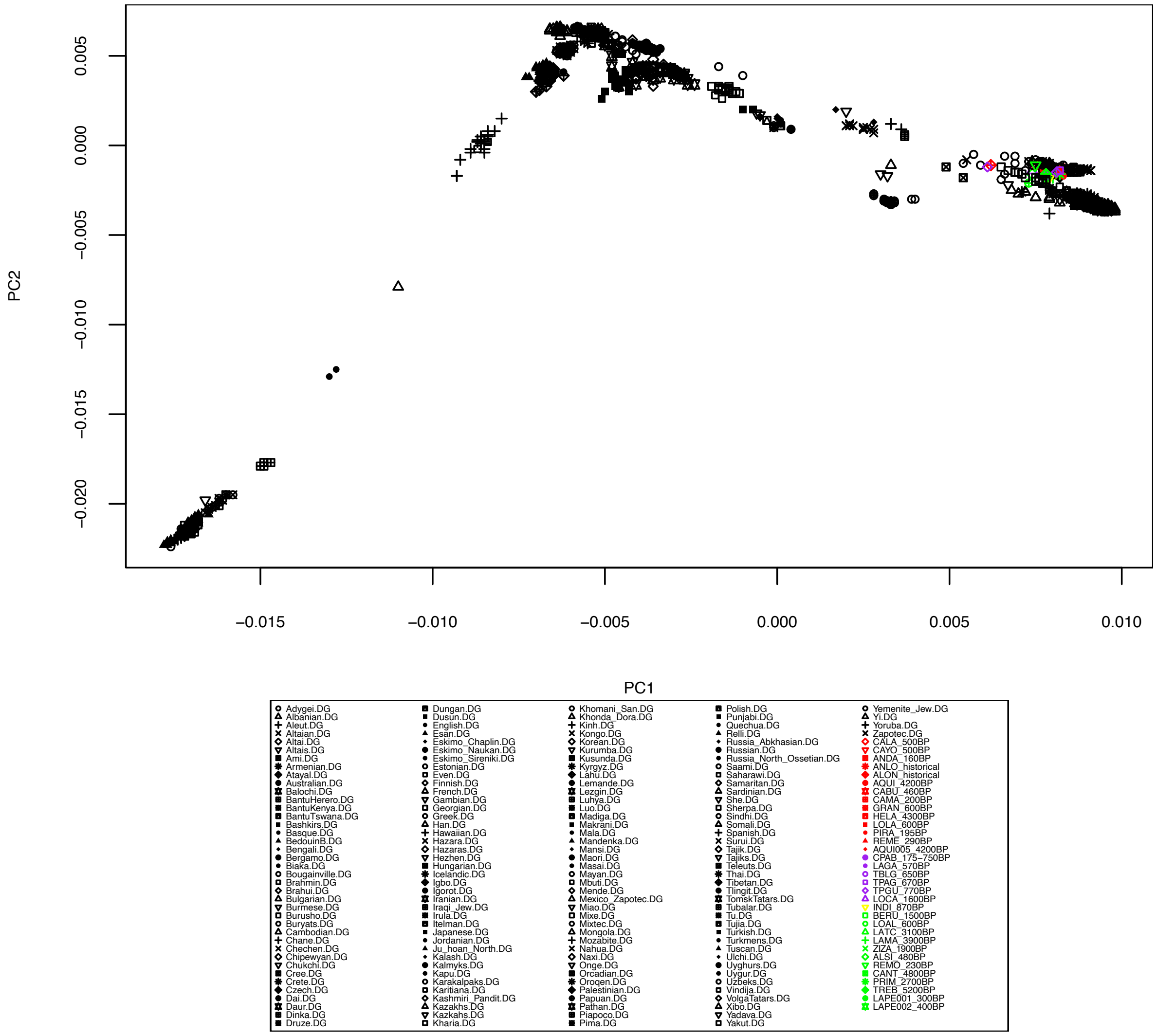


**Figure S1:** Principle Component Analysis. The genetic variation was built with worldwide modern-day individuals genotyped for the 1240K dataset (Mallick et al., 2016) onto which ancient individuals were projected.

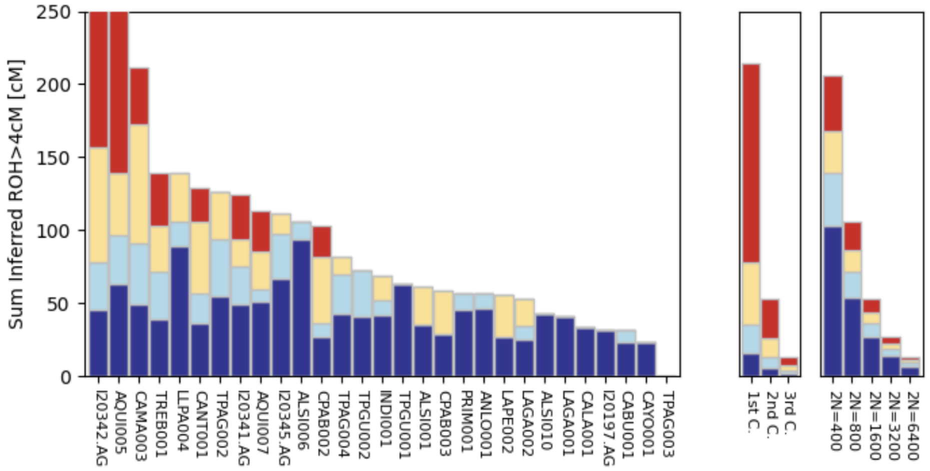


**Figure S2**: Runs of Homozygosity (ROH). ROHs have been assigned for individuals with a SNP coverage above 400,000 on the 1240K with hapROH (Ringbauer et al. 2021).

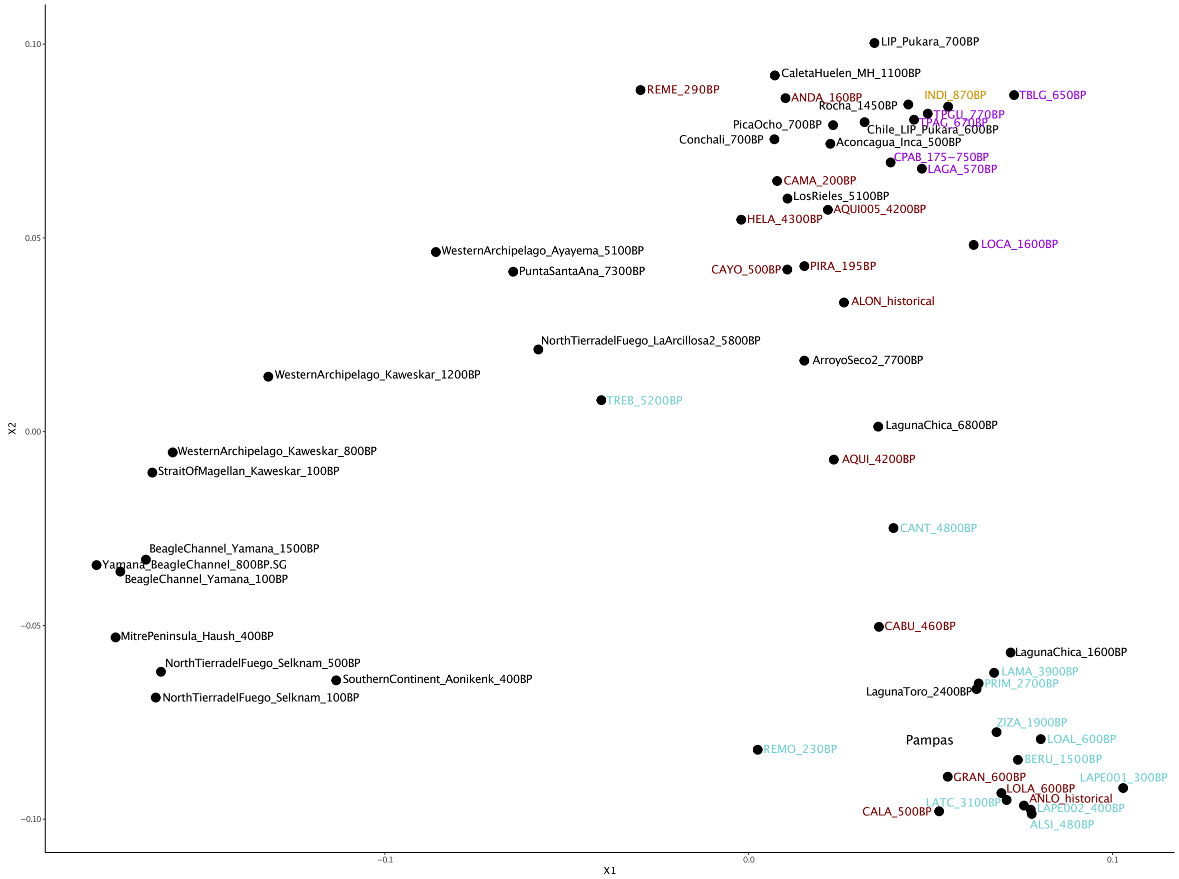


**Figure S3:** Multidimensional Scaling Plot (MDS). The plot is based on 1-$f_3$-outgroup statistics and includes representative individuals/populations of main ancestries identified in the Southern Cone. Newly generated individuals are colored as followed: Northwest Patagonia (red), central and southern Pampas (green), Paraná River Delta and Lower Uruguay River (purple) and eastern lowlands of Uruguay (yellow).

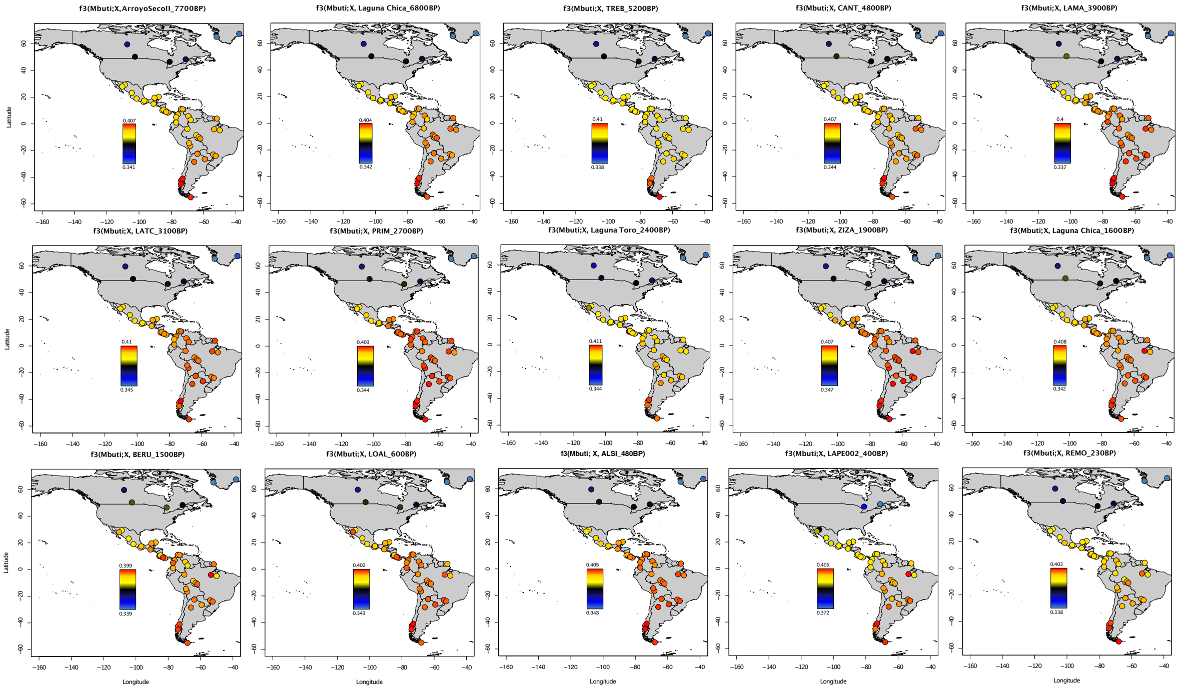


**Figure S4**: F3-outgroup statistics. This analysis measures the shared genetic drift of ancient central and southern Pampas individuals with present-day Indigenous Americans genotyped on the masked Illumina dataset (Reich et al. 2012).

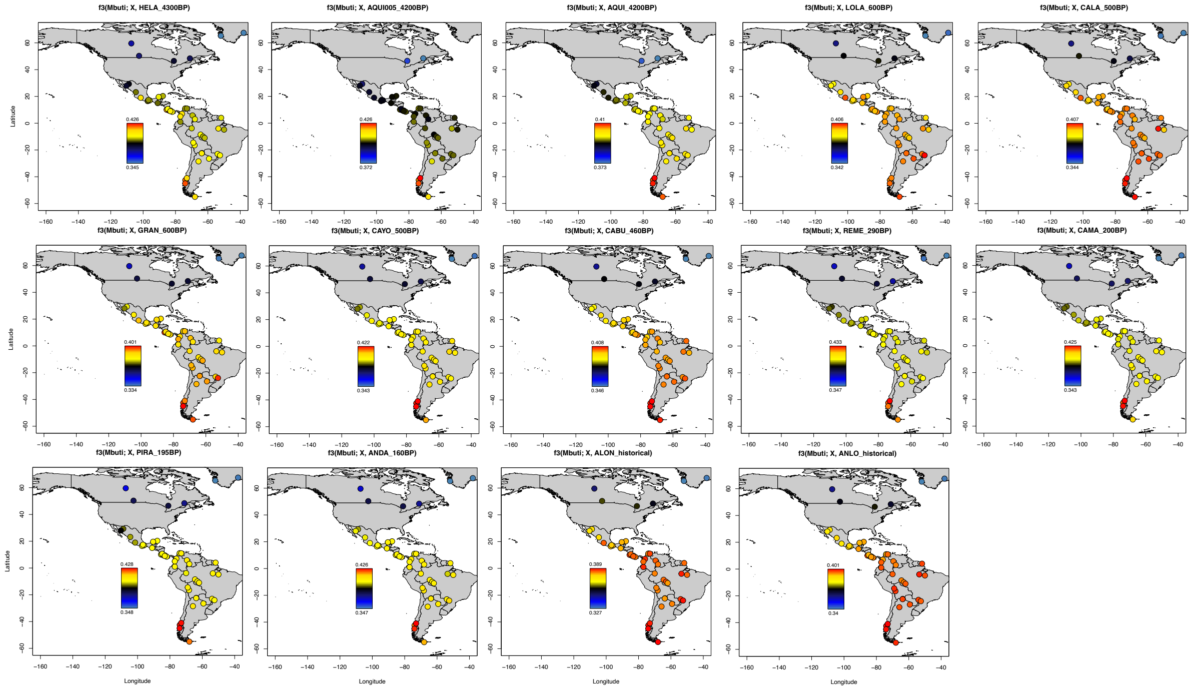


**Figure S5:** F3-outgroup statistics (related to Figure 3A). This analysis measures the shared genetic drift of ancient Northwest Patagonia individuals with present-day Indigenous Americans genotyped on the masked Illumina dataset (Reich et al. 2012).

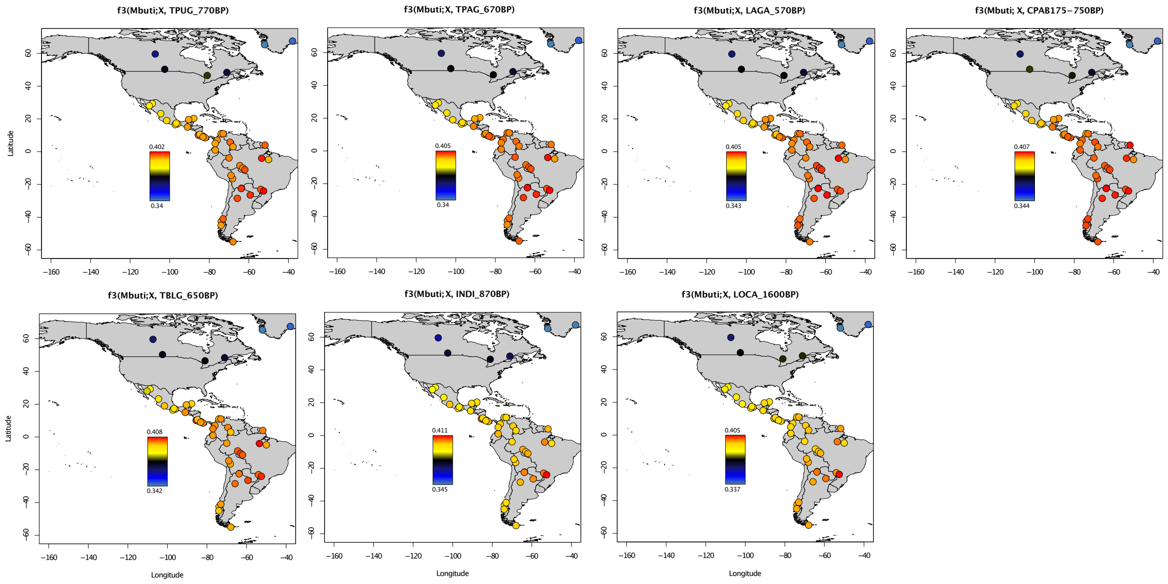


**Figure S6:** $F_3$-outgroup statistics (related to Figure 3A). This analysis measures the shared genetic drift of ancient individuals from Paraná River Delta and Lower Uruguay River, and eastern lowlands of Uruguay, with with present-day Indigenous Americans genotyped on the masked Illumina dataset (Reich et al. 2012).
